## Supplementary Material for "The gene expression landscape of the human locus coeruleus revealed by single-nucleus and spatially-resolved transcriptomics"

August 31, 2023

### Supplementary Tables

**Supplementary Table 1: Summary of experimental design, sample information, and donor demographic details.** Information includes the types of assays performed, donor demographic details, sample IDs, number of Visium tissue areas per sample, and sequencing summary statistics for each sample. The table is provided as an .xlsx file.

**Supplementary Table 2: Differential expression (DE) testing results in pseudobulked Visium SRT data.** Columns include gene ID, gene name, mean log-transformed normalized counts (logcounts) in manually annotated LC and non-LC regions ("mean_logcounts_LC" and "mean_logcounts_nonLC"), $\log_2$ fold change ($\log_2$FC), p-value, false discovery rate (FDR), and columns identifying significant (FDR < 0.05 and FC > 2) and highly significant (FDR < $10^{-3}$ and FC > 3) genes. The table is provided as a .csv file.

**Supplementary Table 3: Spatially variable genes (SVGs) in Visium SRT data.** Results for SVGs identified using nnSVG in Visium SRT data. Columns include gene ID, gene name, overall rank of SVGs identified in replicated tissue areas ("replicated_overall_rank", i.e. top LC-associated SVGs; see **Results**), overall rank of identified SVGs according to average rank across tissue areas ("overall_rank", i.e. including choroid plexus-associated SVGs from one donor; see **Results**), average rank of identified SVGs across individual tissue areas ("average_rank"), number of times (tissue areas) identified within top 100 SVGs ("n_withinTop100"), and ranks within each individual tissue area. The table is provided as a .csv file.

**Supplementary Table 4: Differential expression (DE) testing results for NE neuron cluster vs. other neuronal clusters in snRNA-seq data.** DE testing results comparing NE neuron cluster against other neuronal clusters in snRNA-seq data. Columns include gene ID, gene name, sum of UMI counts across all nuclei ("sum_gene"), average log-transformed normalized counts (logcounts) within the NE neuron cluster ("self_average"), average of average logcounts within other neuronal clusters ("other_average"), combined p-value, false discovery rate (FDR), summary $\log_2$ fold change in the pairwise comparison with the lowest p-value ("summary_logFC"), and column identifying significant (FDR < 0.05 and FC > 2) genes. The table is provided as a .csv file.

**Supplementary Table 5: Differential expression (DE) testing results for NE neuron cluster vs. all other clusters in snRNA-seq data.** DE testing results comparing NE neuron cluster against all other clusters in snRNA-seq data. Columns include gene ID, gene name, sum of UMI counts across all nuclei ("sum_gene"), average log-transformed normalized counts (logcounts) within the NE neuron cluster ("self_average"), average of average logcounts within all other clusters ("other_average"), combined p-value, false discovery rate (FDR), summary $\log_2$ fold change in the pairwise comparison with the lowest p-value ("summary_logFC"), and column identifying significant (FDR < 0.05 and FC > 2) genes. The table is provided as a .csv file.

**Supplementary Table 6: Differential expression (DE) testing results for overlapping set of DE genes in Visium SRT data and snRNA-seq data.** DE testing results for overlapping set of statistically significant (FDR < 0.05 and FC > 2) DE genes in pseudobulked Visium SRT data and between NE neuron cluster vs. all other clusters in snRNA-seq data. Genes are ordered by FC in Visium SRT data. Columns as in Supplementary Tables 2 and 5. The table is provided as a .csv file.

**Supplementary Table 7: Differential expression (DE) testing results for 5-HT neuron cluster in snRNA-seq data.** DE testing results comparing 5-HT neuron cluster against all other neuronal clusters in snRNA-seq data. Columns include gene ID, gene name, sum of UMI counts across all nuclei (“sum_gene”), average log-transformed normalized counts (logcounts) within the 5-HT neuron cluster (“self_average”), average of average logcounts within all other neuronal clusters (“other_average”), combined p-value, false discovery rate (FDR), summary $\log_2$ fold change in the pairwise comparison with the lowest p-value (“summary_logFC”), and column identifying significant (FDR < 0.05 and FC > 2) genes. The table is provided as a .csv file.

### Supplementary Figures

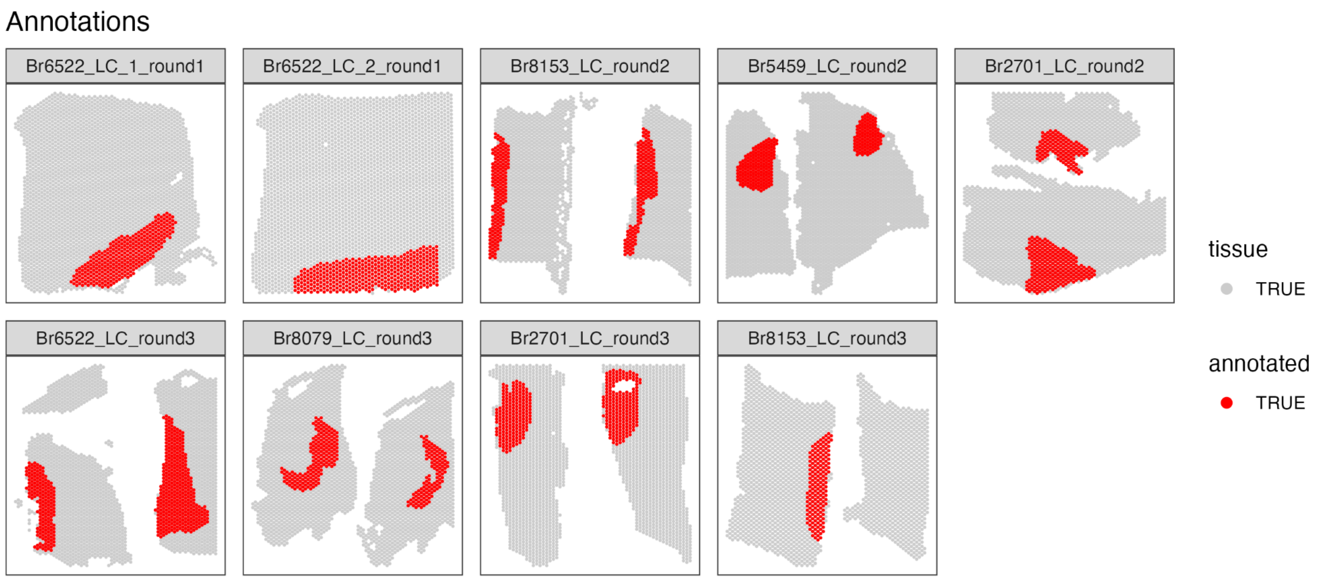


**Supplementary Figure 1: Spot-plot visualizations of manually annotated Visium spots within regions identified as containing LC-NE neurons in SRT data.** For each of the *N*=9 Visium capture areas (hereafter referred to as samples), the spots were manually annotated as being within the LC regions (red) or within the non-LC regions (gray) based on spots containing NE neurons, which were identified by pigmentation, cell size, and morphology on the H&E stained histology images.

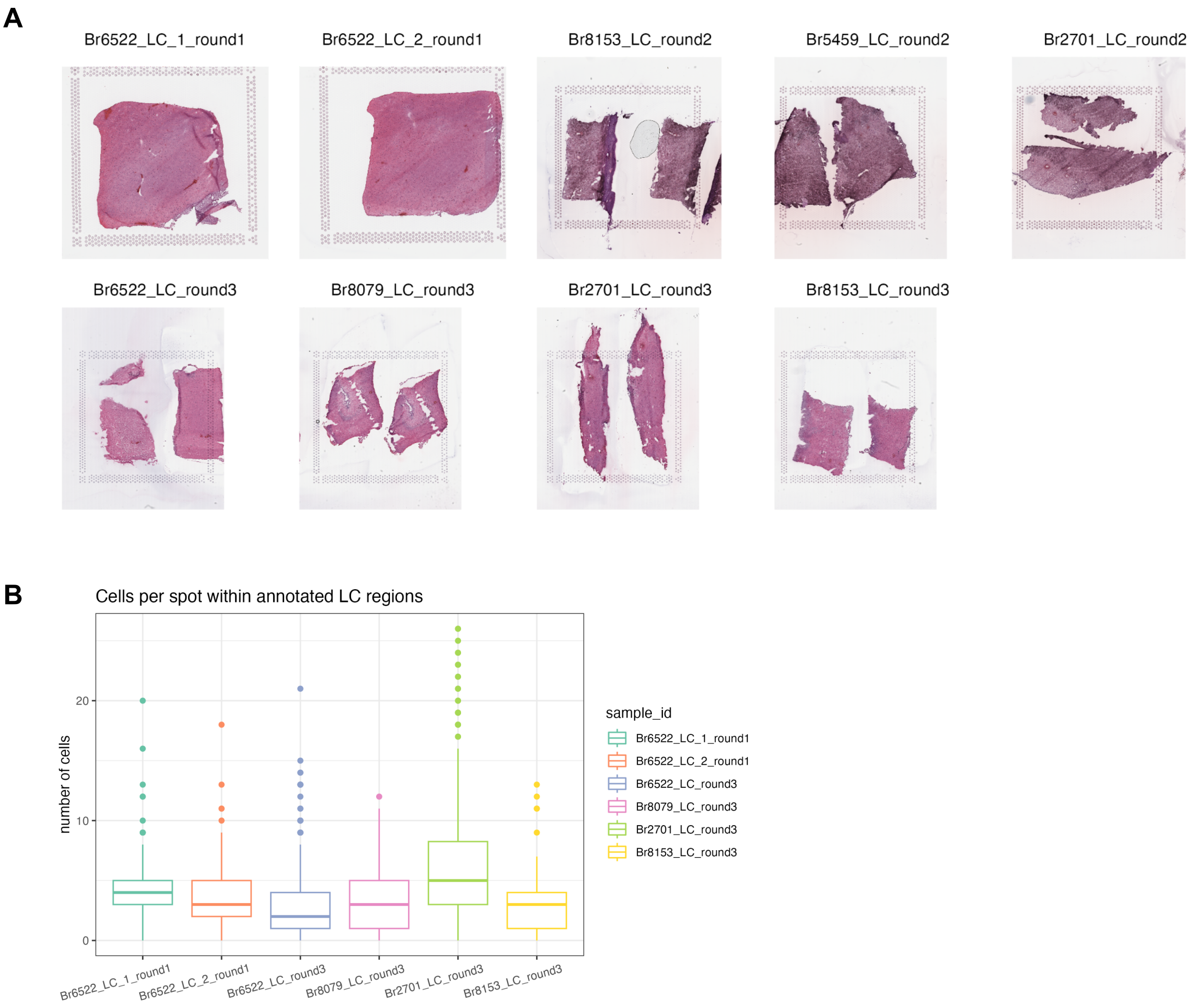


**Supplementary Figure 2: H&E stained histology images and number of cells per spot for SRT data. (A)** Hematoxylin and eosin (H&E) stained histology images for the *N*=9 Visium samples. Higher-resolution images can also be viewed through the Shiny web app (https://libd.shinyapps.io/locus-c_Visium/). **(B)** Estimated number of cells per spot within annotated LC regions in 6 Visium samples, based on application of cell segmentation software (VistoSeg [71]). Boxplots show medians, first and third quartiles, whiskers extending to the furthest values no more than 1.5 times the interquartile range from each quartile, and outliers.

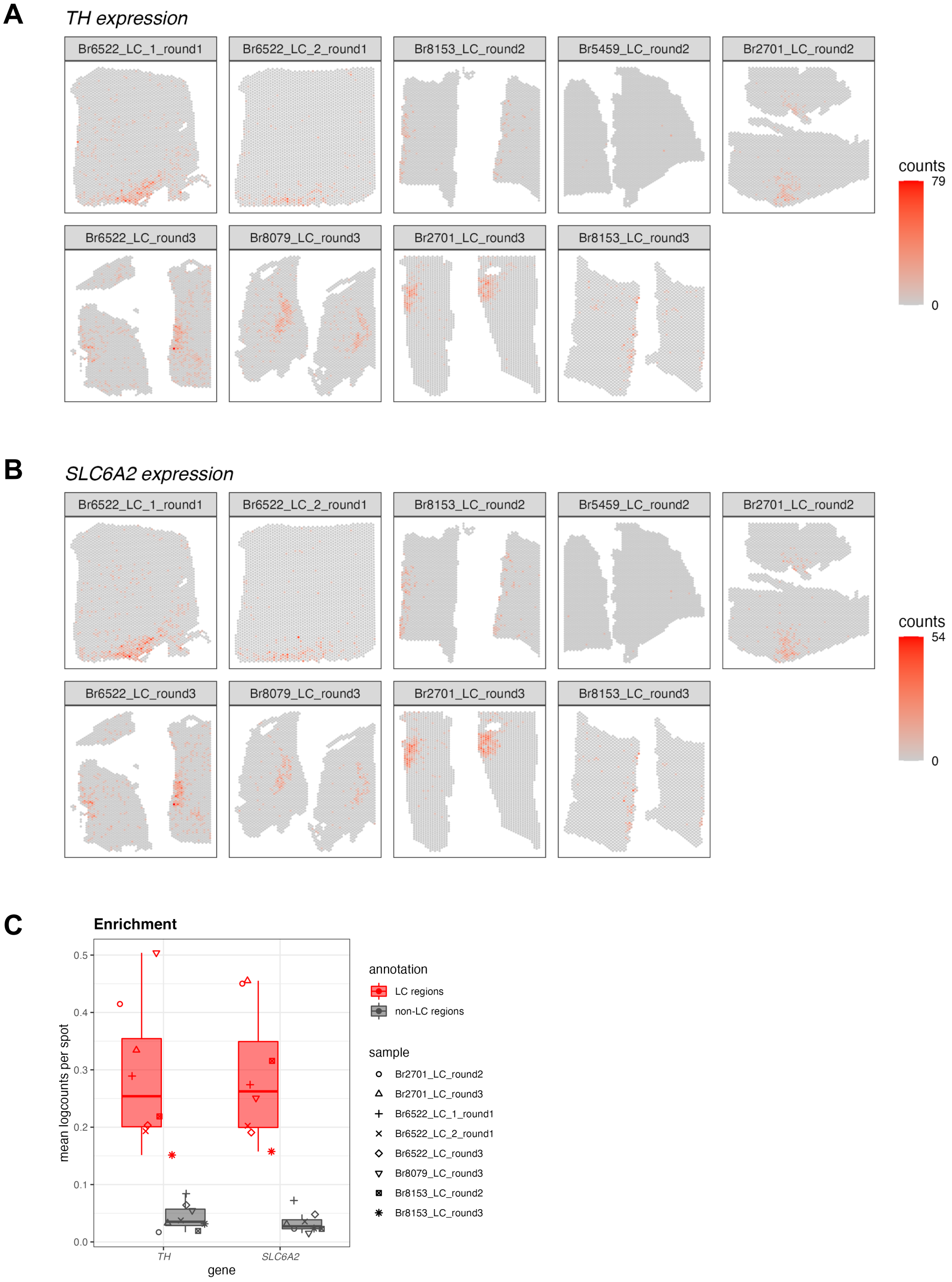


**Supplementary Figure 3: Spatial expression of two NE neuron-specific marker genes in Visium samples for quality control (QC) in SRT data. (A-B)** Spot-plot visualizations of NE neuron marker gene expression (*TH* and *SLC6A2*, **A** and **B**, respectively) in the *N*=9 Visium samples. Color scale shows UMI counts per spot. One sample (Br5459_LC_round2) did not show clear expression of the NE neuron marker genes. This sample was excluded from subsequent analyses, leaving *N*=8 Visium capture areas (samples) from 4 out of the 5 donors. The annotated LC regions are shown in **Supplementary Figure 1**. **(C)** Enrichment of NE neuron marker gene expression (*TH* and *SLC6A2*) within manually annotated LC regions compared to non-LC regions in the *N*=8 Visium samples that passed QC. Boxplots show values as mean log-transformed normalized counts (logcounts) per spot within each region per sample, with samples represented by shapes.

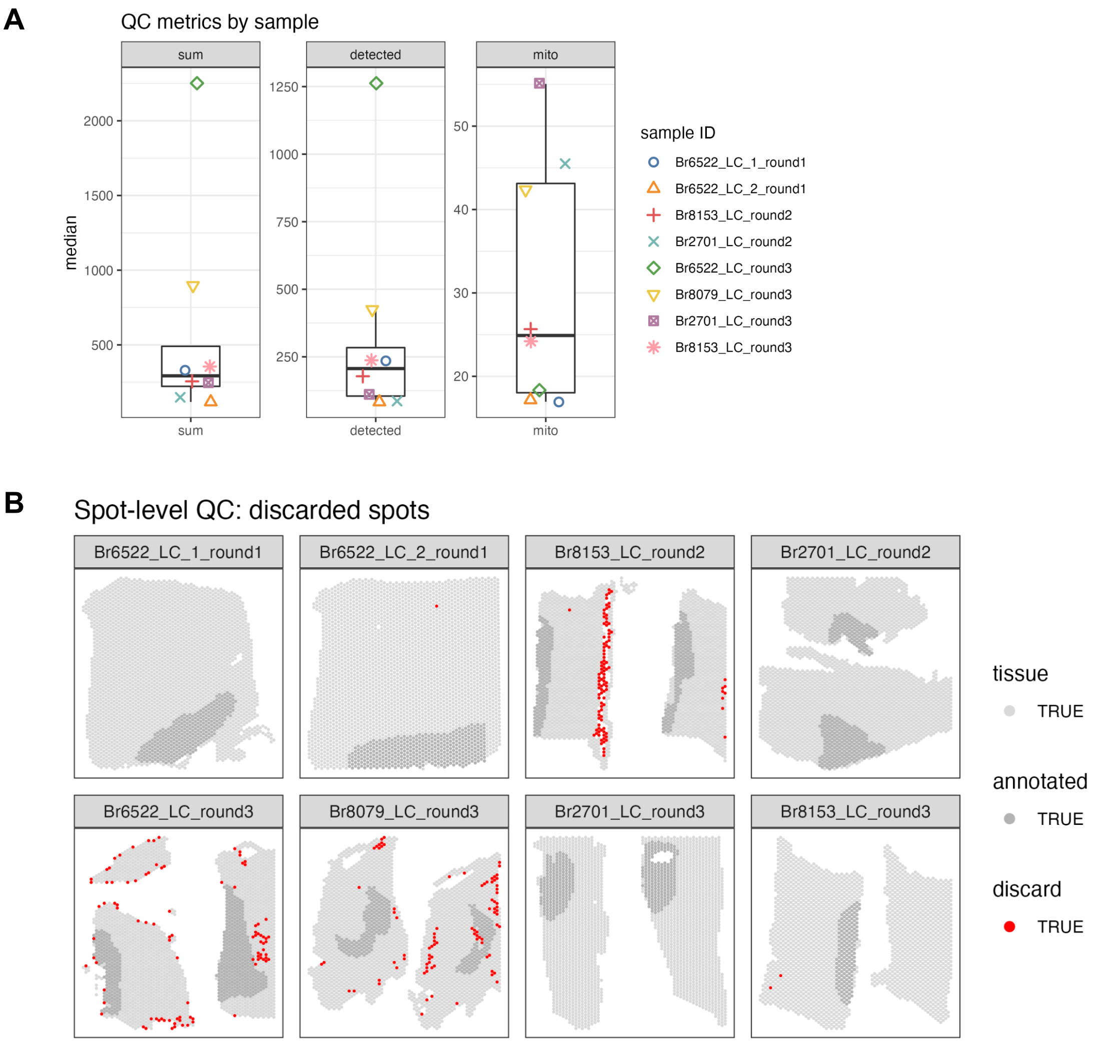


**Supplementary Figure 4: Spot-level quality control (QC) data visualizations for Visium samples in SRT data.** (**A**) QC metrics, medians per sample (from left to right: sum of UMI counts per spot, number of detected genes per spot, and proportion of mitochondrial reads per spot). Boxplots show median for each QC metric per sample, with samples represented by shapes. (**B**) Applying thresholds of 3 median absolute deviations (MADs) to the sum of UMI counts and number of detected genes for each sample identified a total of 287 low-quality spots (red) (1.4% out of 20,667 total spots), which were removed from subsequent analyses. We did not use the proportion of mitochondrial reads for spot-level QC filtering (see **Methods** for more details).

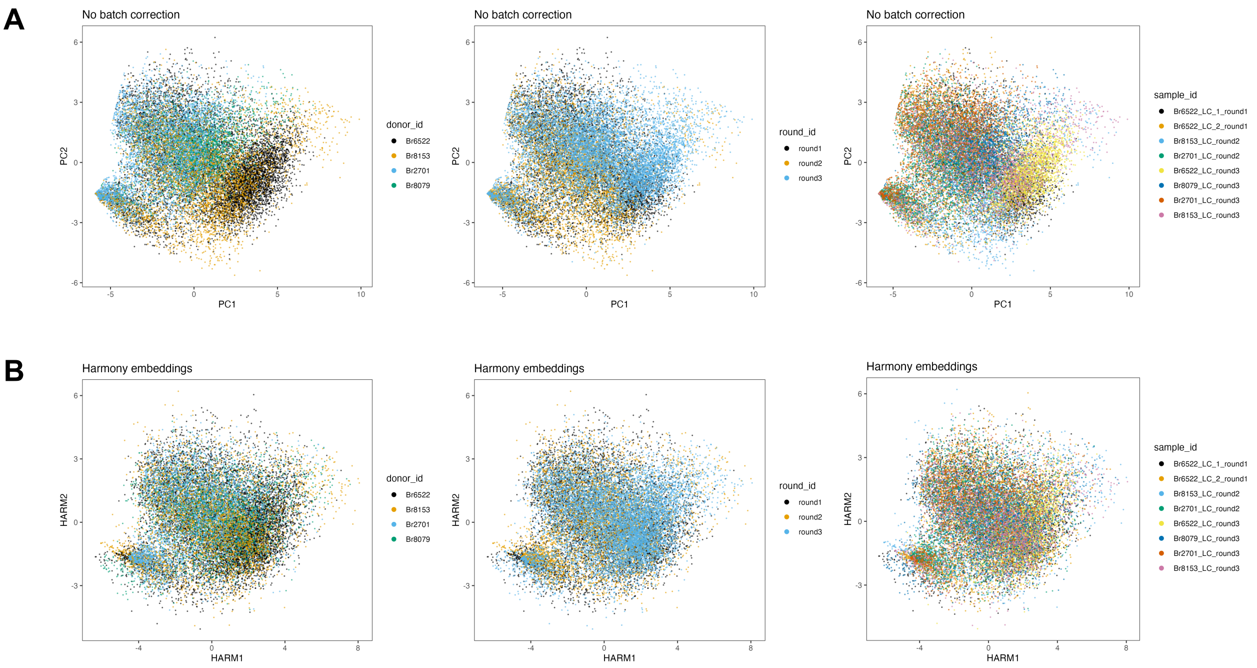


**Supplementary Figure 5: Dimensionality reduction embeddings before and after batch integration across Visium samples in SRT data.** We applied a batch integration tool (Harmony [41]) to remove technical variation in the molecular measurements between the *N*=8 Visium samples from 4 donors. The integrated measurements were subsequently used as the input for spatially-aware clustering using BayesSpace [40]. (**A**) Principal component analysis (PCA) (top 2 PCs) calculated on molecular expression measurements, with spots labeled (left to right) by donor ID, round ID, and sample ID, without applying any batch integration. (**B**) Harmony embeddings (top 2 Harmony embedding dimensions) after applying Harmony batch integration on sample IDs, with spots labeled (left to right) by donor ID, round ID, and sample ID, demonstrating that the technical variation has been reduced.

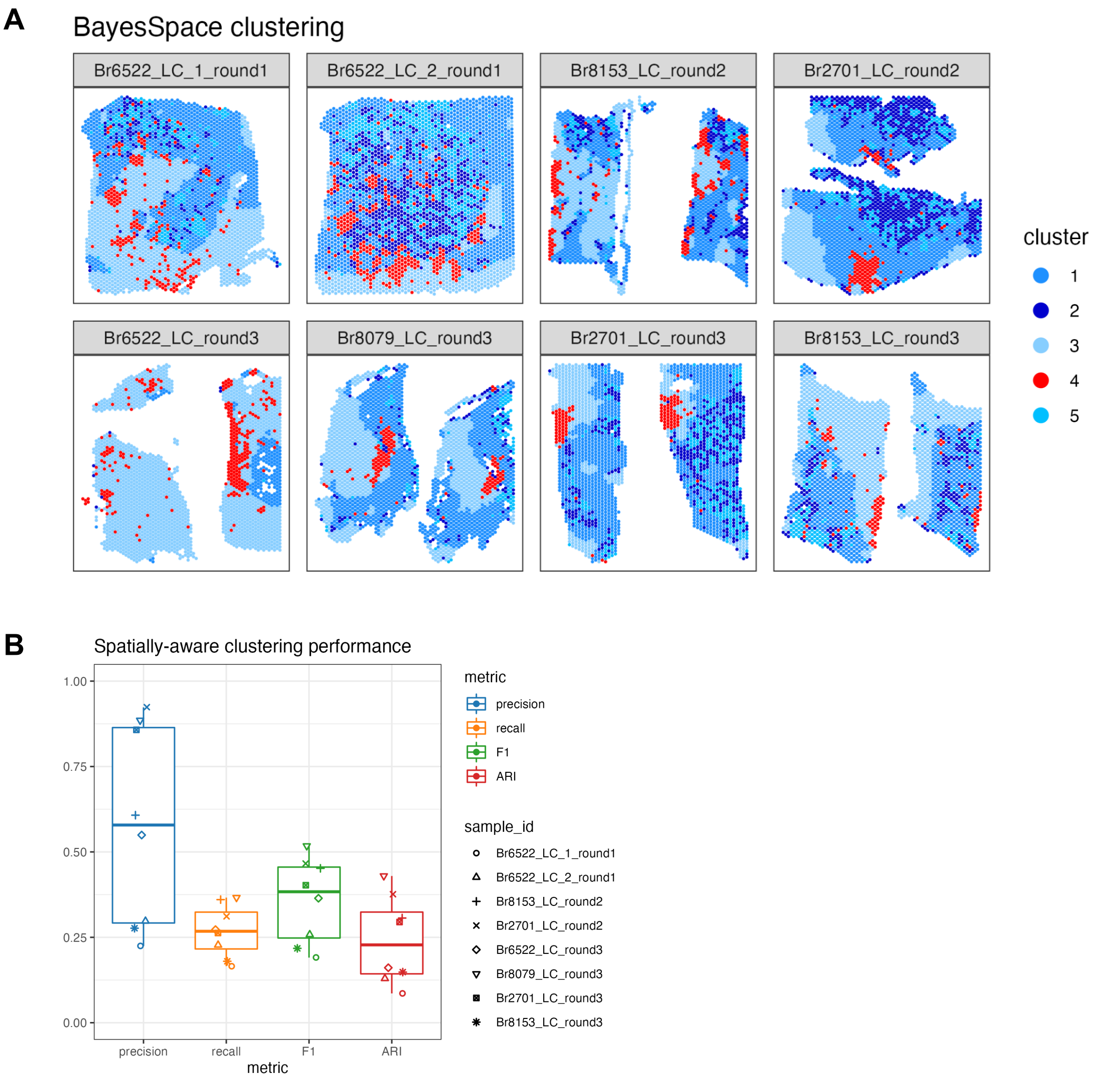


**Supplementary Figure 6: Identifying LC and non-LC regions in a data-driven manner by spatially-aware unsupervised clustering in SRT data.** We applied a spatially-aware unsupervised clustering algorithm (BayesSpace [40]) to investigate whether the LC and non-LC regions in each Visium sample could be annotated in a data-driven manner. (**A**) Using BayesSpace with *k*=5 clusters, we clustered spots from the *N*=8 Visium samples using the Harmony batch-integrated molecular measurements (clustering performed across samples). Cluster 4 (red) corresponds most closely to the manually annotated LC regions. The annotated LC regions are shown in **Supplementary Figure 1**. (**B**) BayesSpace clustering performance evaluated in terms of concordance between cluster 4 (red) and the manually annotated LC region in each sample. Clustering performance was evaluated in terms of precision, recall, F1 score, and adjusted Rand index (ARI) (see **Methods** for definitions).

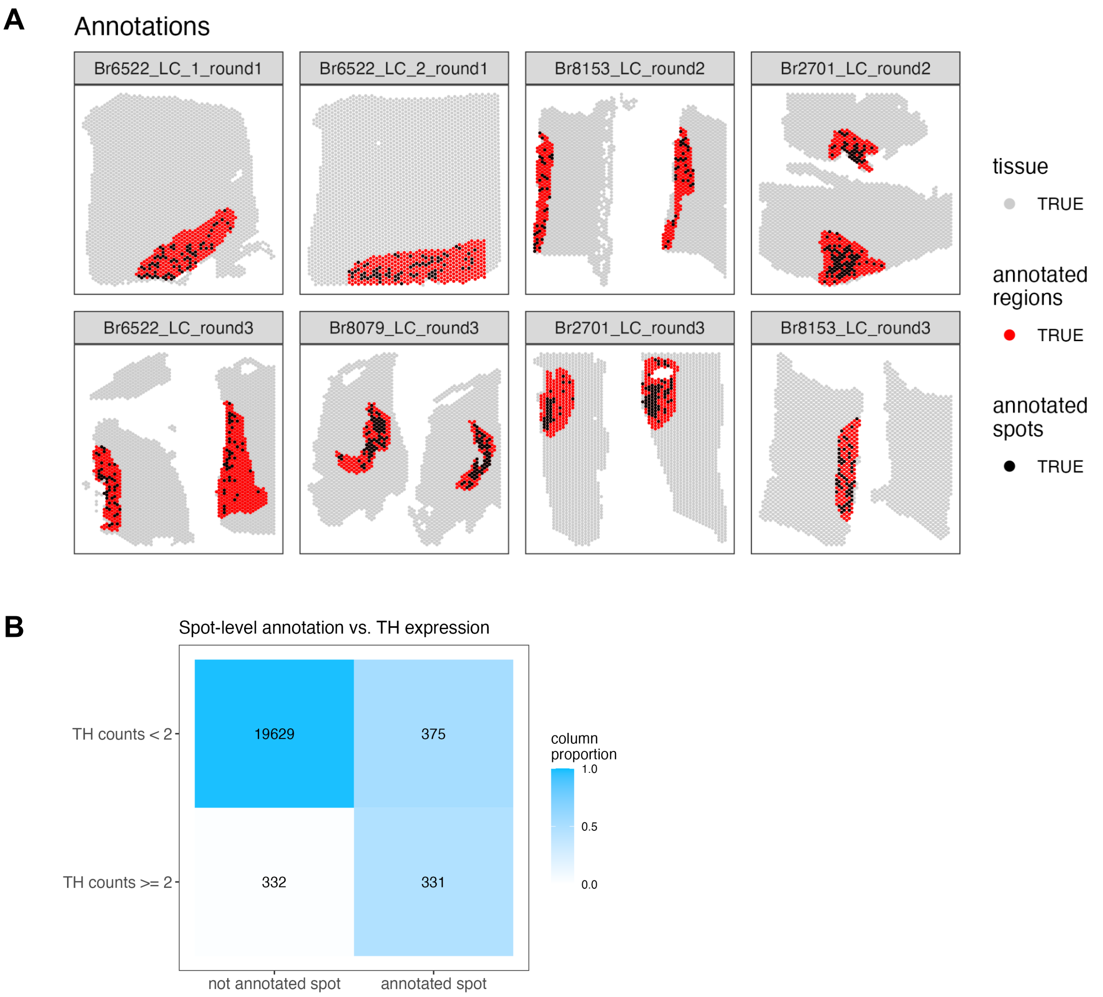


**Supplementary Figure 7: Comparison of spot-level and region-level manual annotations in SRT data. (A)** We manually annotated individual Visium spots (black) overlapping with NE neuron cell bodies within the previously manually annotated LC regions (red), based on pigmentation, cell size, and morphology from the H&E stained histology images, in the *N*=8 Visium samples. (**B**) We observed relatively low overlap between spots with expression of the NE neuron marker gene *TH* (>=2 observed UMI counts per spot) and the set of annotated individual spots. The differences included both false positives (annotated spots that were not *TH*+) and false negatives (*TH*+ spots that were not annotated). Therefore, we did not use the spot-level annotations for subsequent analyses, and instead used the LC region-level annotations for all further analyses.

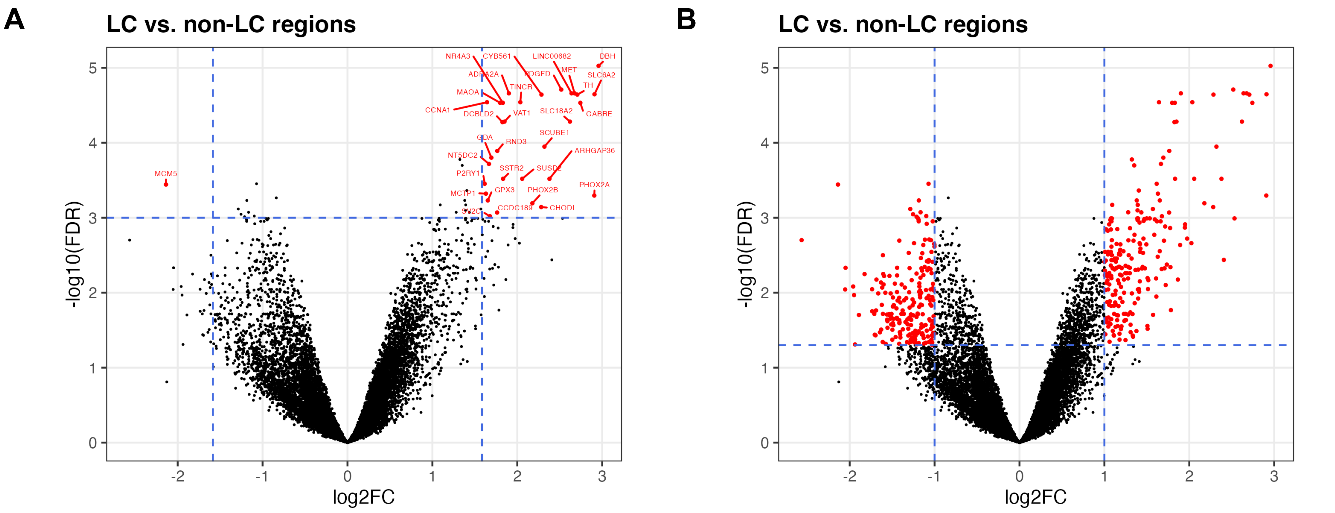


**Supplementary Figure 8: Results from differential expression (DE) analysis to identify expressed genes associated with LC regions in SRT data.** We performed DE testing between the manually annotated LC and non-LC regions by pseudobulking spots, defined as aggregating UMI counts from the combined set of spots, within the annotated LC and non-LC regions in each sample. (**A**) Using a false discovery rate (FDR) significance threshold of $10^{-3}$ and an expression fold-change (FC) threshold of 3 (dashed blue lines), we identified 32 highly significant genes (red points). (**B**) Using standard significance thresholds of FDR < 0.05 and expression FC > 2, we identified 437 significant genes (red). Vertical axes are on reversed $\log_{10}$ scale, and horizontal axes are on $\log_2$ scale. Additional details are provided in **Supplementary Table 2**.

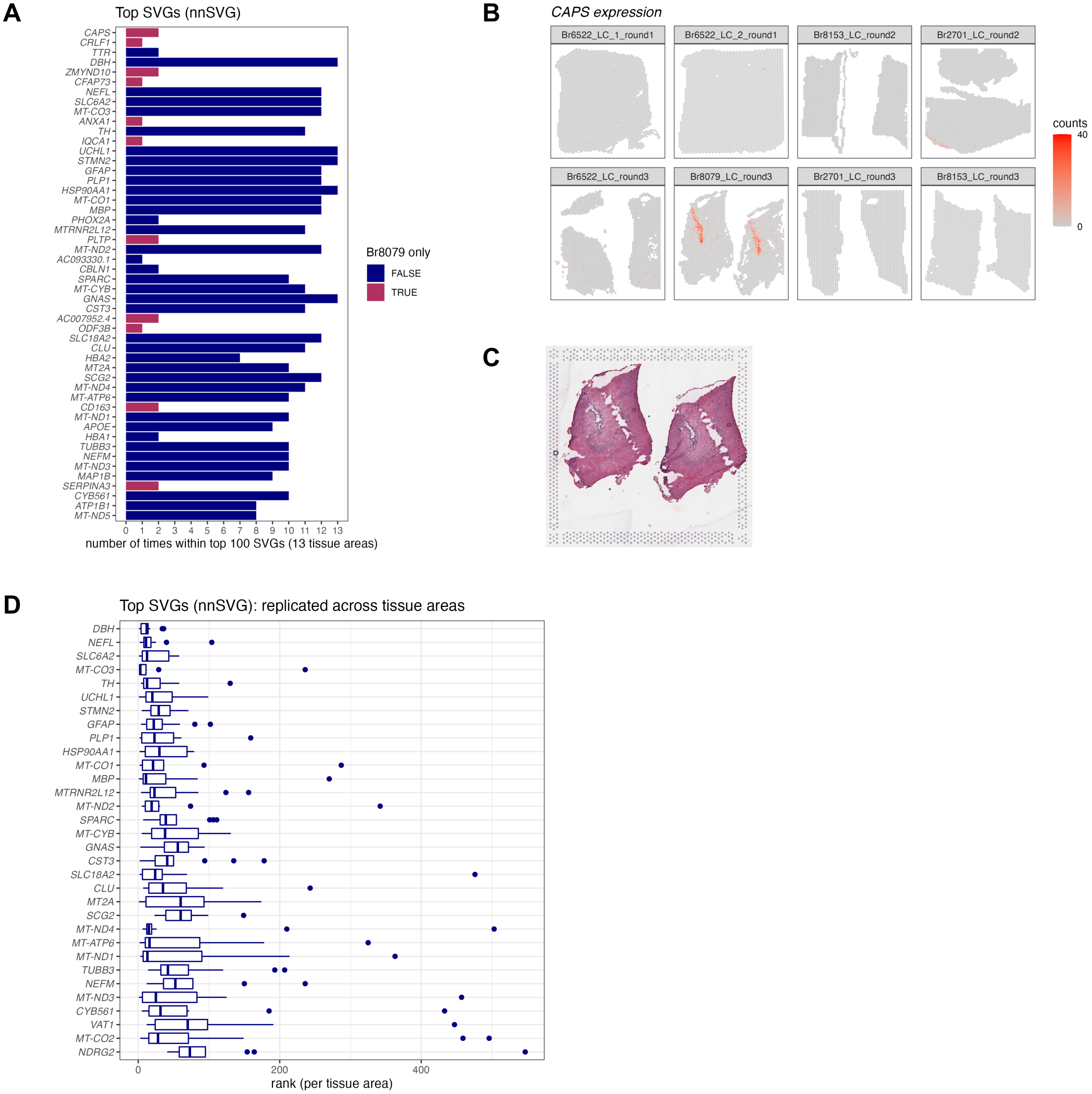


**Supplementary Figure 9: Results from applying nnSVG to identify spatially variable genes (SVGs) in SRT data.** We applied nnSVG [43], a method to identify spatially variable genes (SVGs), in the Visium SRT samples. We ran nnSVG within each contiguous tissue area containing a manually annotated LC region (13 tissue areas in the *N*=8 Visium samples) and calculated an overall ranking of top SVGs by averaging the ranks per gene from each tissue area. **(A)** The top 50 ranked SVGs from this analysis included a subset (11 out of 50) of genes that were highly ranked in samples from only one donor (Br8079, genes highlighted in maroon). We determined that this was due to the inclusion of a section of the choroid plexus adjacent to the LC for this donor. Bars show the number of times (out of 13 tissue areas) each gene was included within the top 100 SVGs. Rows are ordered by overall average ranking in descending order. **(B)** Spatial expression of *CAPS*, a choroid plexus marker gene, in the *N*=8 Visium samples. **(C)** Histology image showing the two tissue areas for sample Br8079_LC_round3. **(D)** In order to focus on LC-associated SVGs, we calculated an overall average ranking of SVGs that were each included within the top 100 SVGs in at least 10 out of the 13 tissue areas, which identified 32 highly-ranked, replicated LC-associated SVGs. Boxplots show the ranks in each tissue area. Rows are ordered by the overall average ranking in descending order.

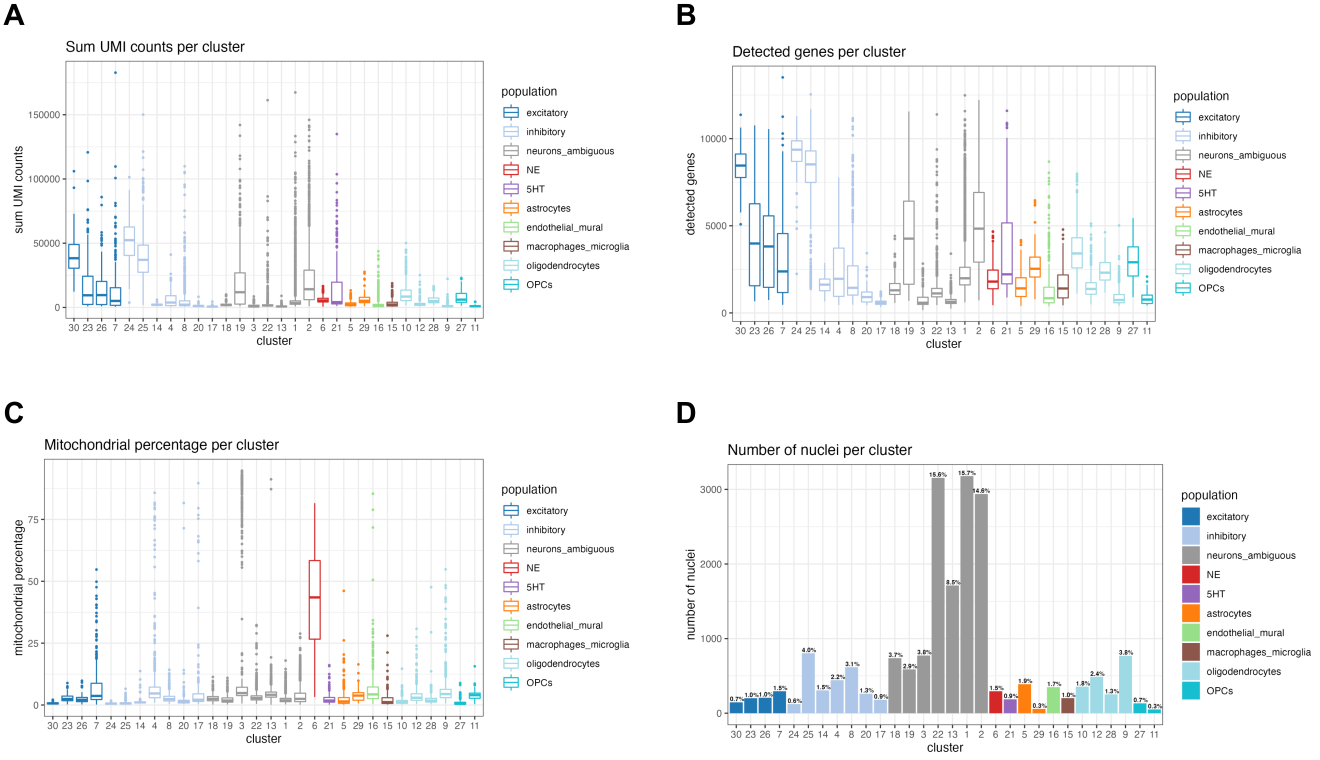


**Supplementary Figure 10: Distribution of nucleus-level quality control (QC) metrics across unsupervised clusters in snRNA-seq data. (A)** Sum of UMI counts per nucleus and cluster, **(B)** total number of detected genes per nucleus and cluster, **(C)** percentage of mitochondrial reads per nucleus and cluster, and **(D)** number and percentage of nuclei per cluster. We observed an unexpectedly high percentage of mitochondrial reads in the NE neuron cluster (cluster 6, red, **C**). Since NE neurons were of particular interest for analysis, we did not remove nuclei with a high percentage of mitochondrial reads during QC filtering.

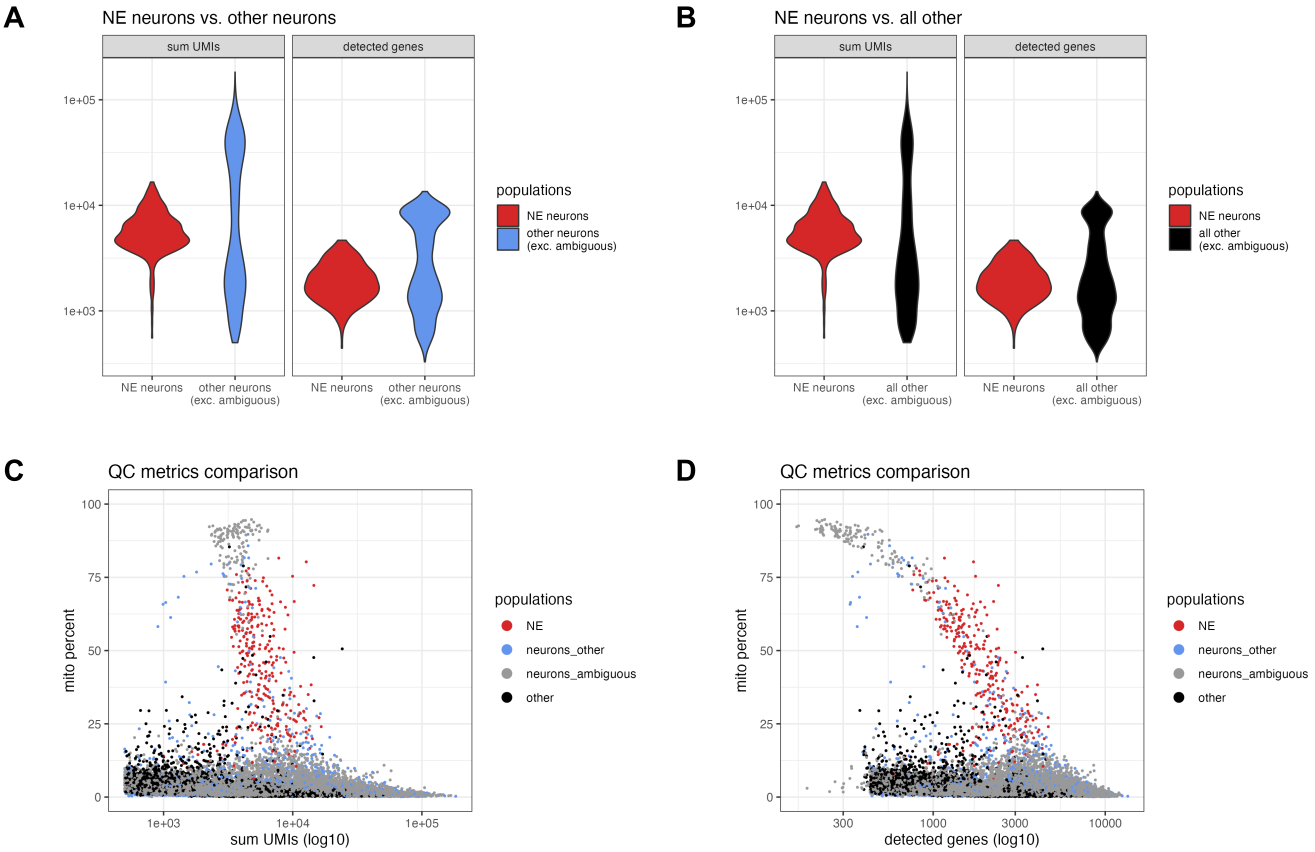


**Supplementary Figure 11: Additional quality control (QC) evaluations for NE neuron cluster in snRNA-seq data.** Additional quality control (QC) comparisons for snRNA-seq data showing **(A)** sum of UMI counts and number of detected genes for the NE neuron cluster compared to other neurons (excluding ambiguous neurons), **(B)** sum of UMI counts and number of detected genes for the NE neuron cluster compared to all other cells (excluding ambiguous neurons), and **(C-D)** percent mitochondrial reads (*y*-axis) vs. **(C)** sum of UMI counts and **(D)** number of detected genes, showing individual nuclei for NE neurons (red), other neurons (blue), ambiguous neurons (gray), and other non-neuronal populations (black). The ambiguous neuron category (gray) includes a distinct group of droplets with the overall highest mitochondrial proportions and lowest number of detected genes, representing likely damaged nuclei and/or debris, which are clearly separated from the NE neurons (red). See also **Supplementary Figure 10** for QC metrics calculated for all clusters.

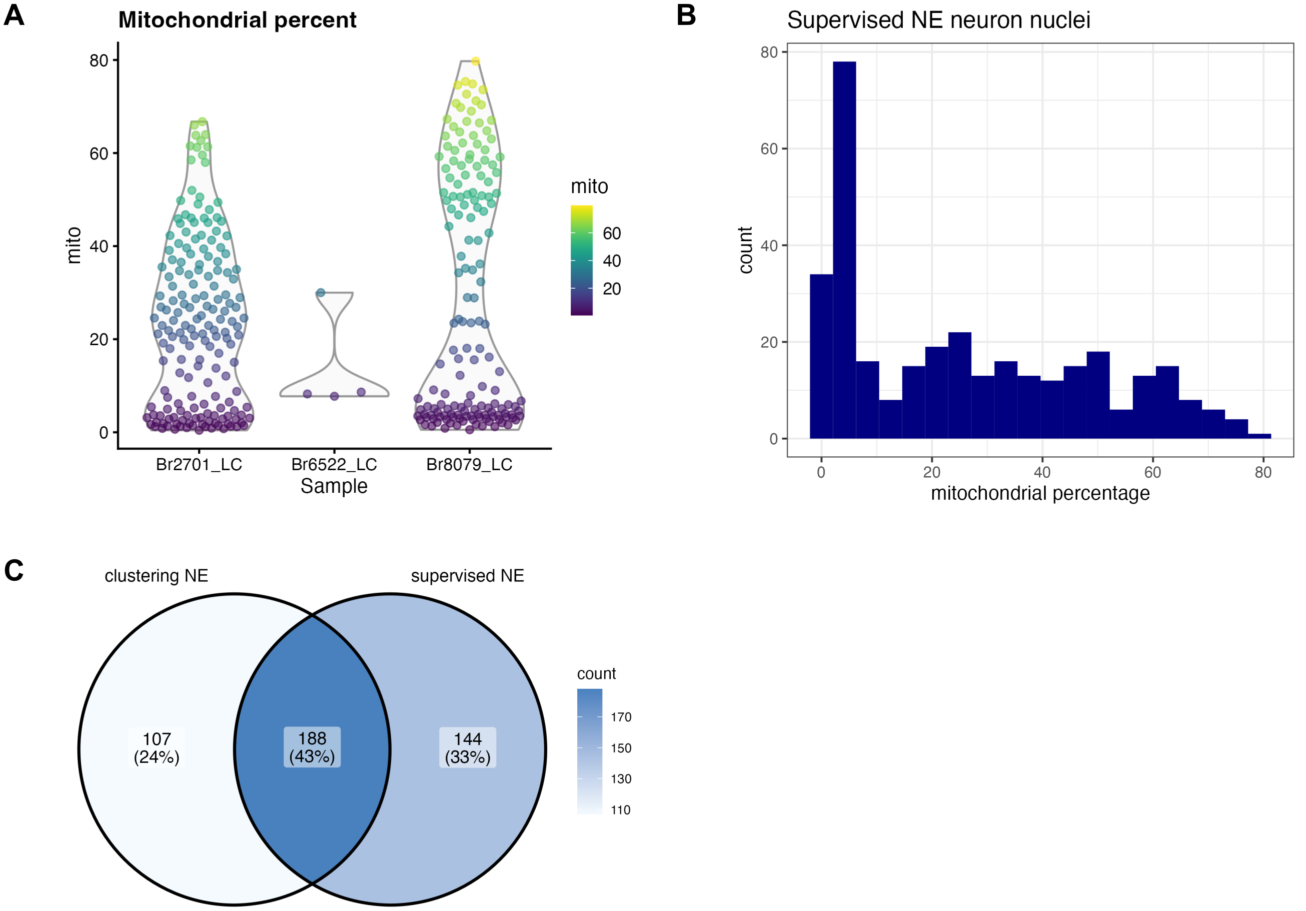


**Supplementary Figure 12: Supervised identification of NE neuron nuclei by thresholding on expression of NE neuron marker genes in snRNA-seq data.** We applied a supervised strategy to identify NE neuron nuclei by simply thresholding on expression of NE neuron marker genes (selecting nuclei with >=1 UMI counts of both *DBH* and *TH*). We observed a higher than expected proportion of mitochondrial reads within this set of nuclei, and did not filter on this parameter during QC processing, in order to retain these nuclei. **(A)** Percentage of mitochondrial reads within the supervised set of nuclei by donor (Br2701, Br6522, and Br8079). **(B)** Histogram showing percentage of mitochondrial reads within the supervised set of nuclei across all donors. **(C)** Venn diagram showing overlap between NE neuron cluster identified by unsupervised clustering (left) and NE neuron population identified by supervised thresholding (right). Values display the number of nuclei.

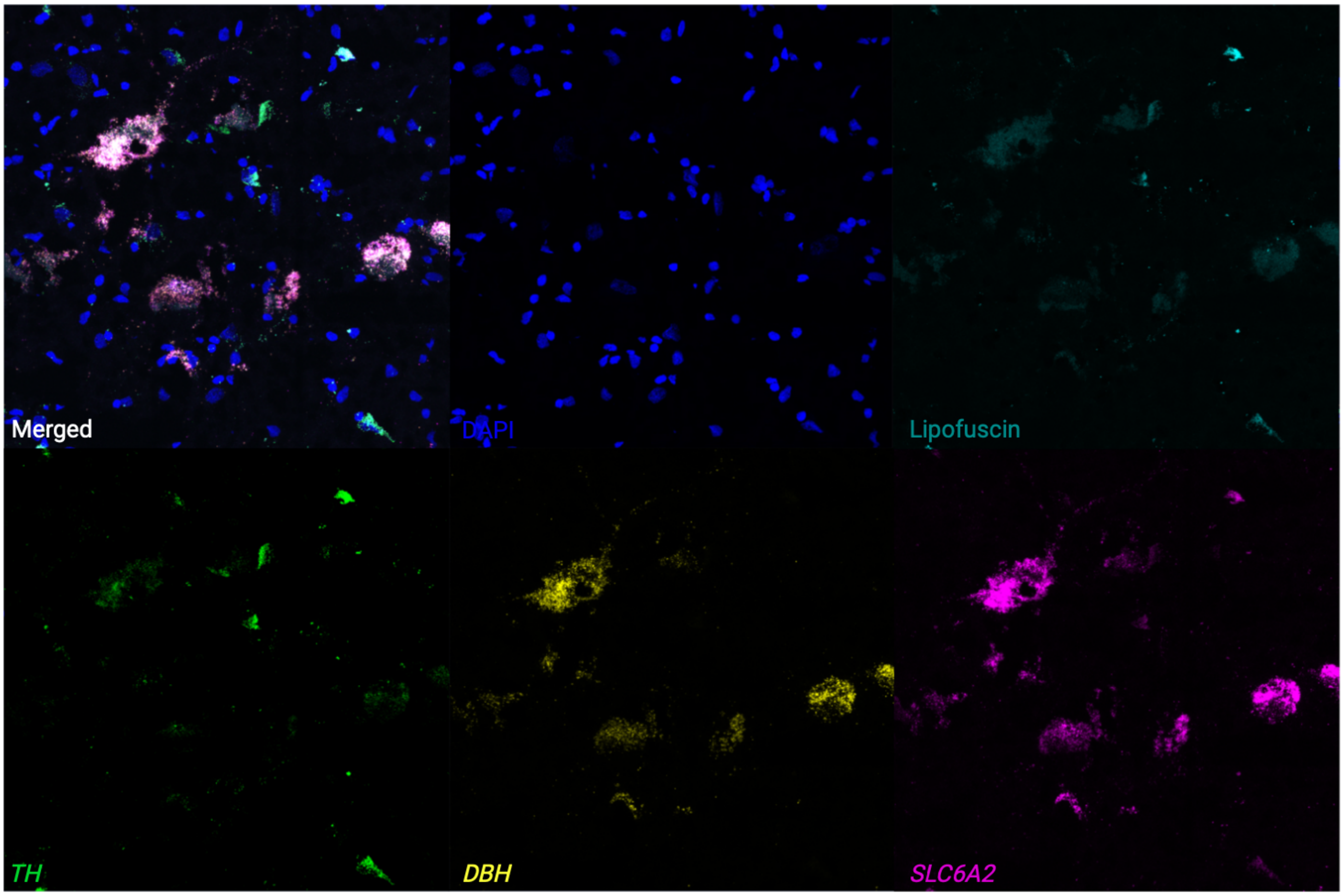


**Supplementary Figure 13: Expression of NE neuron marker genes in individual cells using RNAscope and high-magnification confocal imaging.** We applied RNAscope [37] and high-magnification confocal imaging to visualize expression of NE neuron marker genes (*TH* in green, *DBH* in yellow, and *SLC6A2* in pink, with white representing all three colors overlapping), DAPI stain for nuclei (blue), and lipofuscin (teal) on additional tissue sections from an additional independent donor, Br8689. Top row: merged channels, DAPI, and lipofuscin. Bottom row: individual channels for NE neuron marker genes in separate panels. The figure displays a region within the LC region from a single tissue section, demonstrating clear co-localization of expression of the three NE neuron marker genes (white points in merged channels) within individual cells. Scale bar: 20 μm.

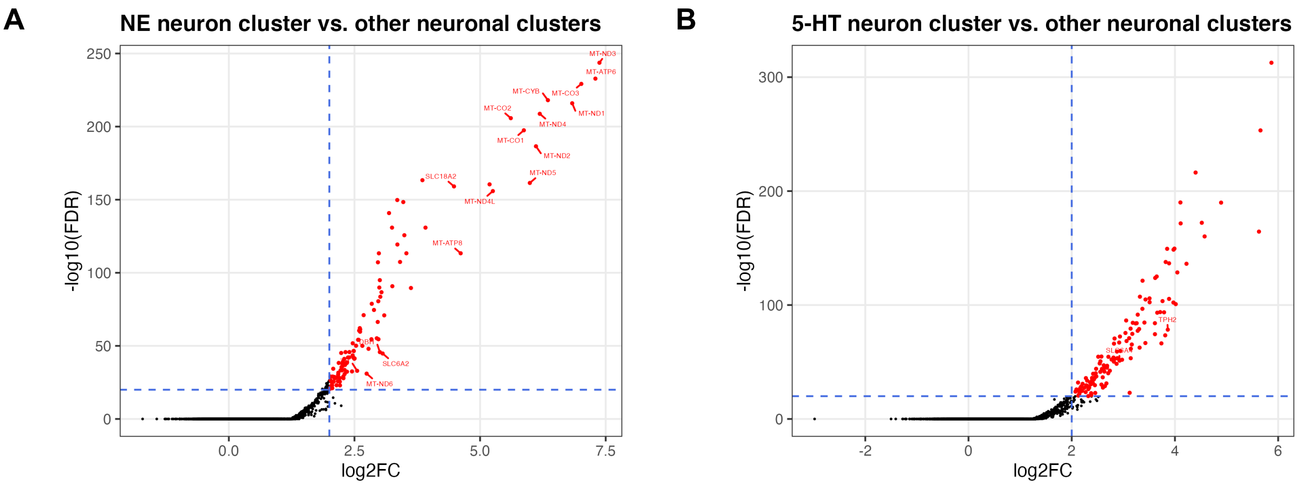


**Supplementary Figure 14: DE testing results between neuronal clusters in the LC and surrounding region in snRNA-seq data. (A)** Volcano plot showing 327 statistically significant DE genes (FDR < 0.05 and FC > 2) elevated in expression within the NE neuron cluster compared to all other neuronal clusters (excluding ambiguous) captured in this region. The significant DE genes include known NE neuron marker genes (*DBH*, *TH*, *SLC6A2*, and *SLC18A2*) and mitochondrial genes. **(B)** Volcano plot showing 361 statistically significant DE genes (FDR < 0.05 and FC > 2) elevated in expression within the 5-HT neuron cluster compared to all other neuronal clusters (excluding ambiguous) captured in this region. The significant DE genes include known 5-HT neuron marker genes (*TPH2* and *SLC6A4*). Vertical axes are on reversed $\log_{10}$ scale, and horizontal axes are on $\log_2$ scale. Additional details are provided in **Supplementary Tables 4, 6**.

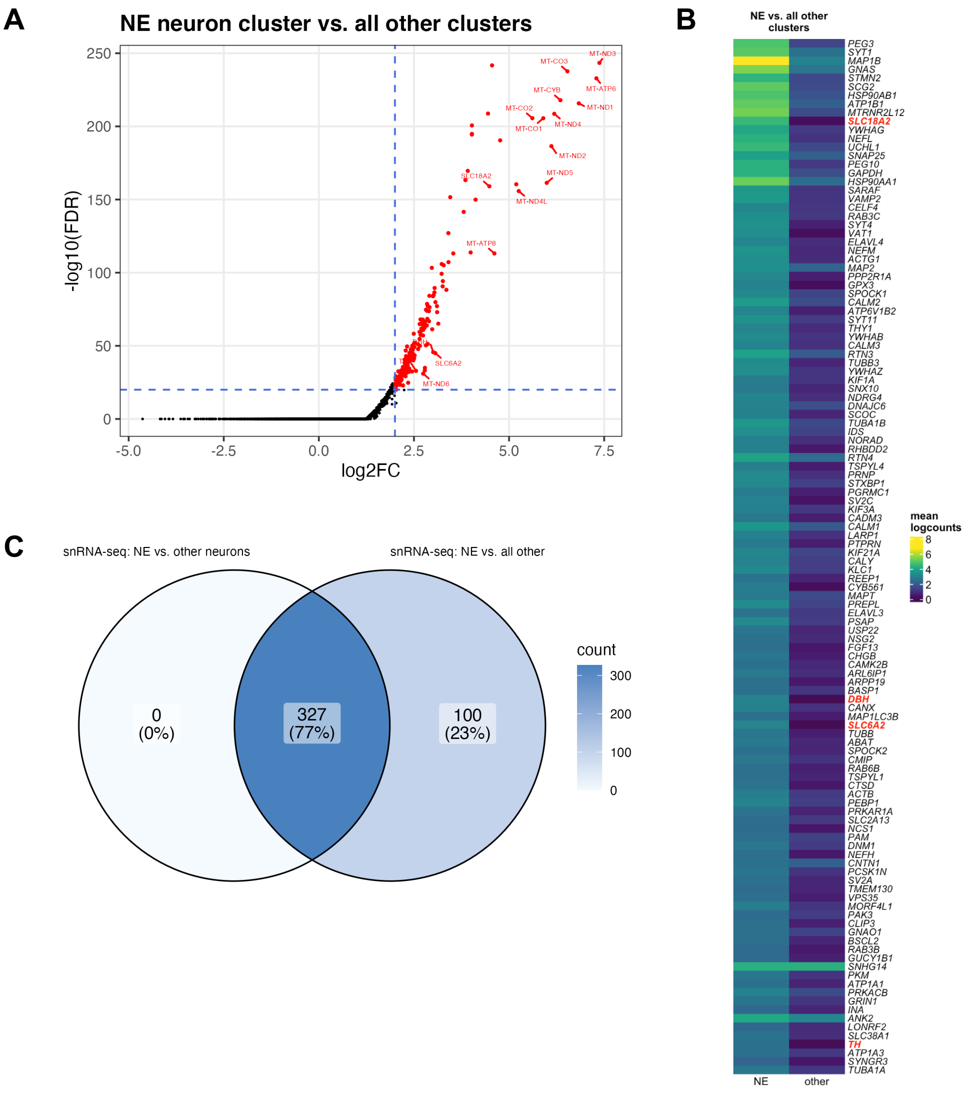


**Supplementary Figure 15: DE testing results between NE neuron cluster and all other clusters in snRNA-seq data. (A)** Volcano plot showing 427 statistically significant DE genes (FDR < 0.05 and FC > 2) elevated in expression within the NE neuron cluster compared to all other clusters (neuronal and non-neuronal, excluding ambiguous neuronal). The significant DE genes include known NE neuron marker genes (*DBH*, *TH*, *SLC6A2*, and *SLC18A2*) and mitochondrial genes. **(B)** Heatmap displaying top 120 genes, ranked in descending order by FDR, excluding mitochondrial genes, with NE neuron marker genes described in text highlighted in red. The full list of 427 genes including mitochondrial genes is provided in **Supplementary Table 5**. Heatmap values represent mean logcounts in the NE neuron cluster and mean logcounts per cluster averaged across all other clusters (excluding ambiguous neuronal). **(C)** Venn diagram showing overlap between sets of statistically significant DE genes identified between NE neuron cluster and other neuronal clusters (left) and between NE neuron cluster and all other clusters (right). Values display the number of genes.

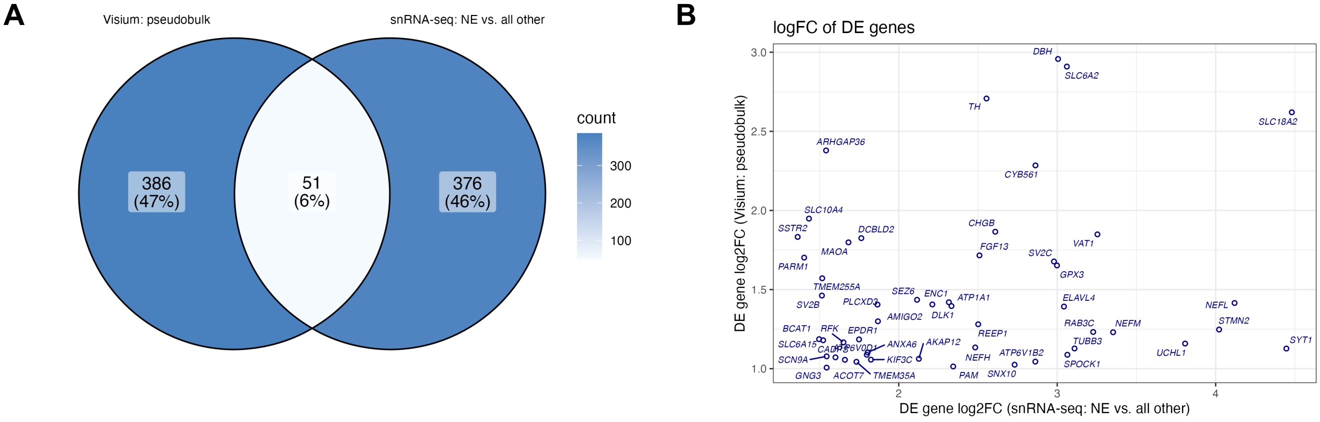


**Supplementary Figure 16: Overlap and comparison between DE genes identified in SRT and snRNA-seq data. (A)** Venn diagram showing overlap between sets of statistically significant DE genes identified in SRT data (pseudobulked LC vs. non-LC regions, left) and snRNA-seq data (NE neuron cluster vs. all other clusters excluding ambiguous neuronal, right). Values display the number of genes. **(B)** Comparison between $\log_2$ fold change (FC) for 51 genes identified as statistically significant DE genes (FDR < 0.05 and FC > 2) in both SRT data (pseudobulked LC vs. non-LC regions, vertical axis) and snRNA-seq data (NE neuron cluster vs. all other clusters excluding ambiguous neuronal, horizontal axis). Additional details for these 51 genes are provided in **Supplementary Table 6**.

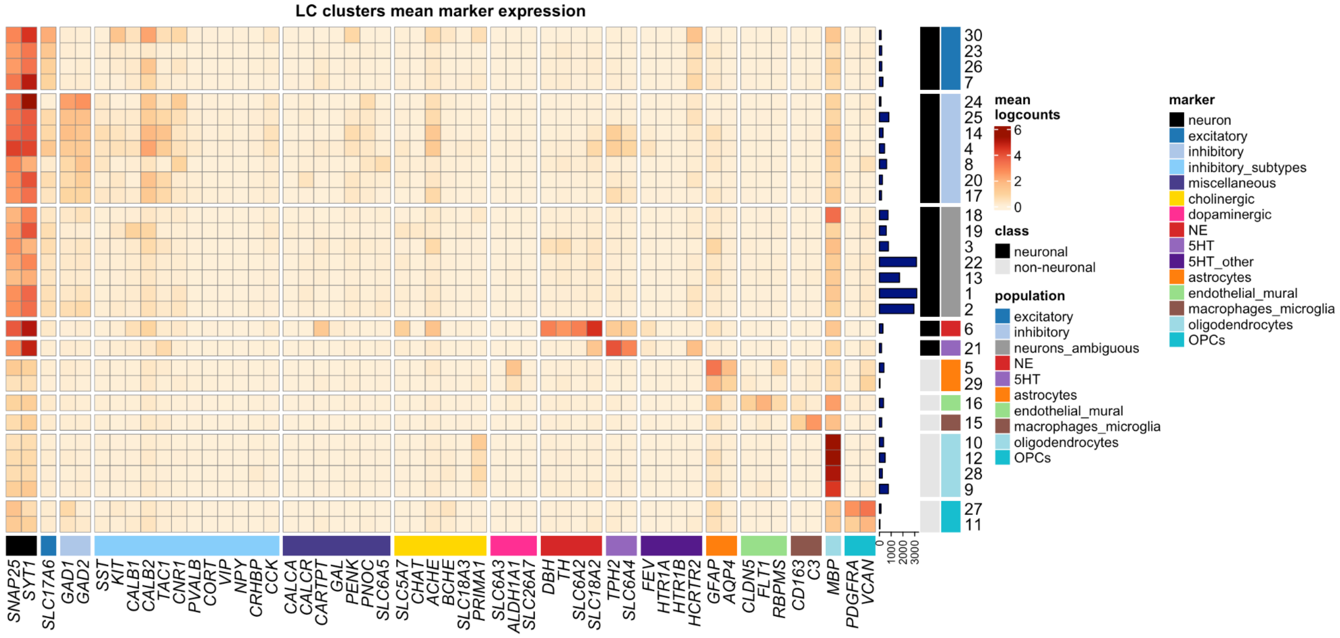


**Supplementary Figure 17: Unsupervised clustering results showing additional inhibitory neuronal, miscellaneous, dopaminergic, and cholinergic marker genes in snRNA-seq data.** Extended form of heatmap displayed in **Figure 3A**, showing additional inhibitory neuronal marker genes (light blue), miscellaneous marker genes including neuropeptides and receptors included for comparison with [32] (dark blue-purple), cholinergic marker genes (yellow), dopaminergic marker genes (pink), and additional 5-HT neuron marker genes for comparison with [45] (dark purple; *SLC6A18* is not shown since we observed zero expression of this gene in the nuclei that passed filtering). We observed diversity in expression of inhibitory neuronal marker genes across inhibitory neuronal subpopulations (additional results in **Supplementary Figure 22**), and we observed expression of cholinergic marker genes within NE neurons (additional results in **Supplementary Figure 26**). We did not observe expression of dopaminergic marker genes within the NE neuron cluster (additional results in **Supplementary Figures 24-25**). Percentages of nuclei per cluster are shown in **Supplementary Figure 10D**.

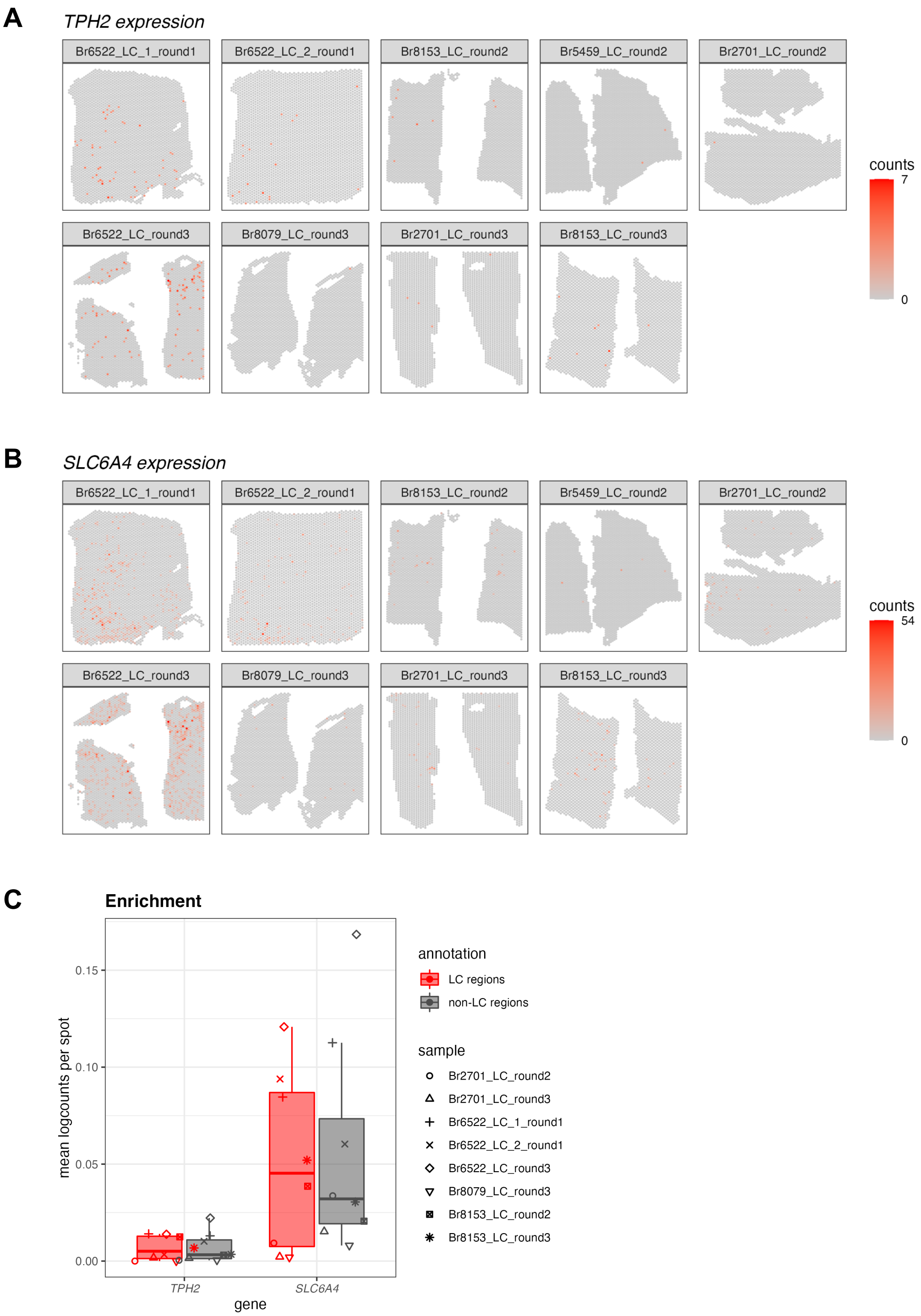


**Supplementary Figure 18: Spatial expression and enrichment analysis of 5-HT neuron marker genes in Visium SRT samples. (A-B)** We visualized the spatial expression of 5-HT (5-hydroxytryptamine or serotonin) neuron marker genes (*TPH2* and *SLC6A4*) in the *N*=9 initial Visium SRT samples within the Visium SRT samples, which showed that the population of 5-HT neurons was distributed across both the LC and non-LC regions. The annotated LC regions are shown in **Supplementary Figure 1**. **(C)** Enrichment of 5-HT neuron marker gene expression (*TPH2* and *SLC6A4*) within manually annotated LC regions compared to non-LC regions in the *N*=8 Visium SRT samples that passed QC (see **Supplementary Figure 3**). Boxplots show values as mean log-transformed normalized counts (logcounts) per spot within each region per sample, with samples represented by shapes.

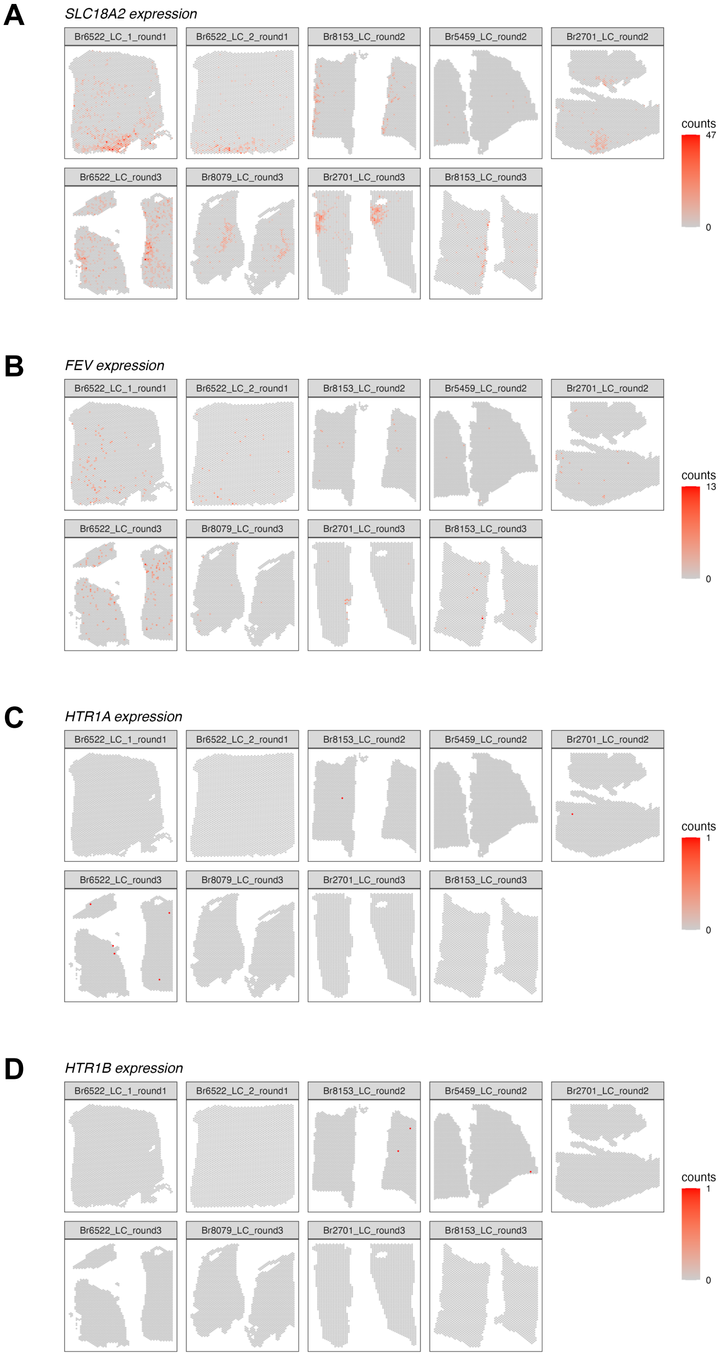


**Supplementary Figure 19: Spatial expression of additional marker genes for 5-HT neurons in Visium SRT samples.** We visualized the spatial expression of **(A-B)** additional marker genes for 5-HT neurons (*SLC18A2*, *FEV*) and **(C-D)** 5-HT autoreceptor genes (*HTR1A*, *HTR1B*) in the *N*=9 initial Visium samples. *SLC6A18* is not shown since we observed zero expression of this gene in the Visium samples. Color scale shows UMI counts per spot.

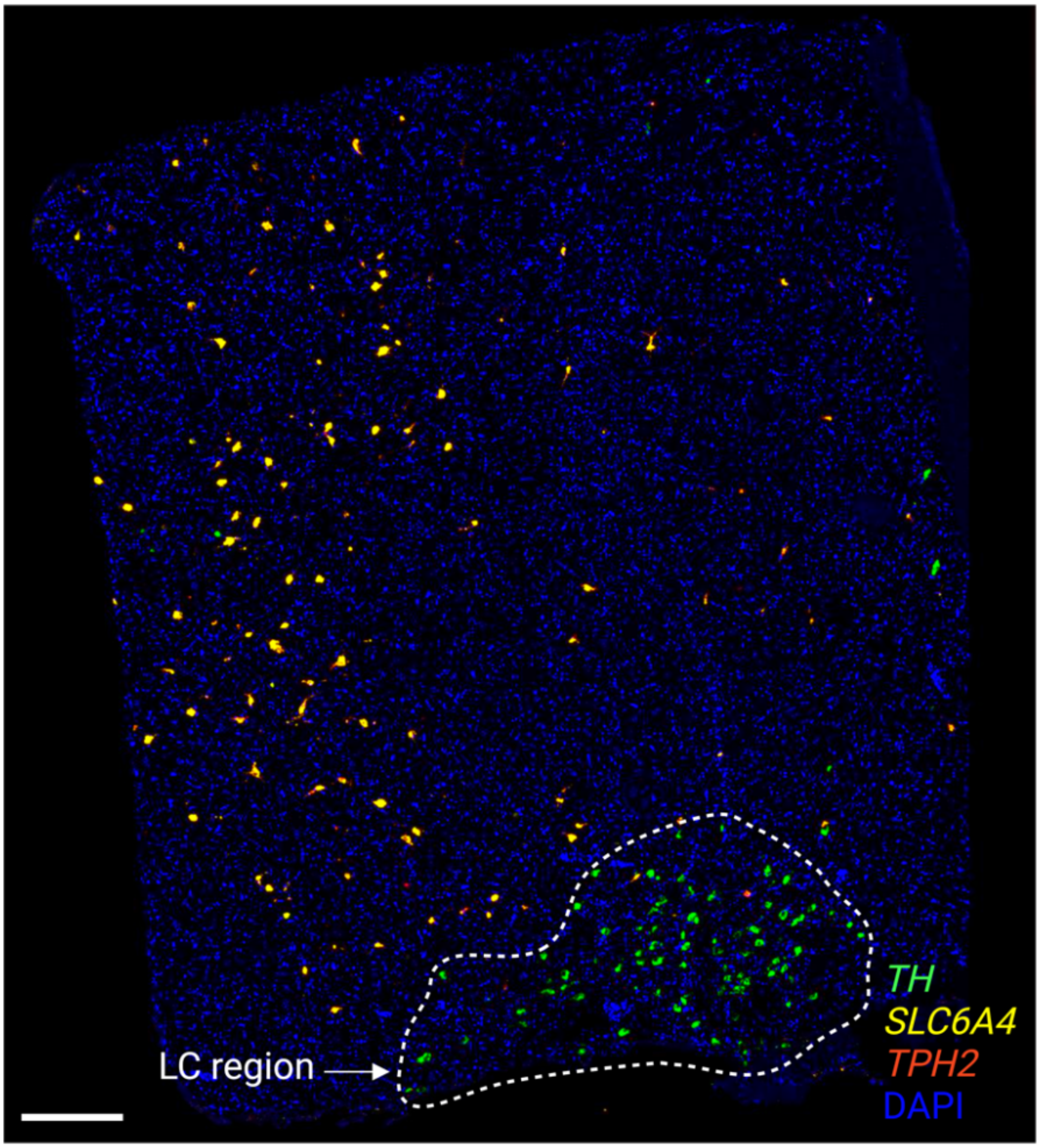


**Supplementary Figure 20: Expression of NE neuron and 5-HT neuron marker genes using RNAscope.** We applied RNAscope [37] to visualize expression of an NE neuron marker gene (*TH*) as well as 5-HT neuron marker genes (*TPH2* and *SLC6A4*) within an additional tissue section from donor Br6522, demonstrating that the NE and 5-HT marker genes were expressed within distinct cells and that the NE and 5-HT neuron populations were not localized within the same regions. Scale bar: 500 μm.

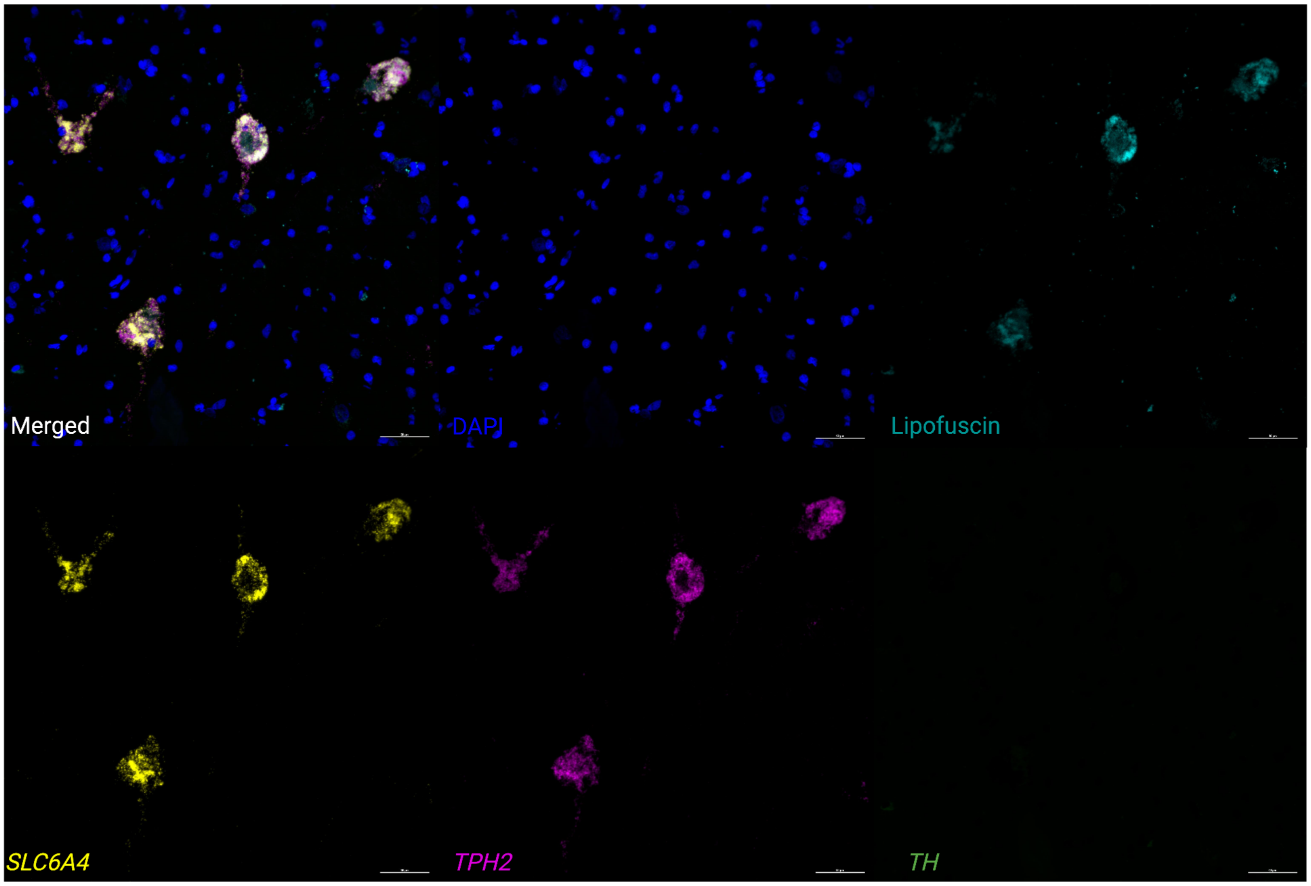


**Supplementary Figure 21: Expression of 5-HT neuron marker genes using RNAscope.** We applied RNAscope [37] and high-magnification confocal imaging to visualize expression of two 5-HT neuron marker genes (*TPH2* and *SLC6A4*) and an NE neuron marker gene (*TH*) within an additional tissue section from donor Br6522, demonstrating that the 5-HT neuron marker genes were co-expressed within individual cells. Image region corresponds to a small region within the RNAscope image displayed in Supplementary Figure 20. Scale bar: 50 µm.

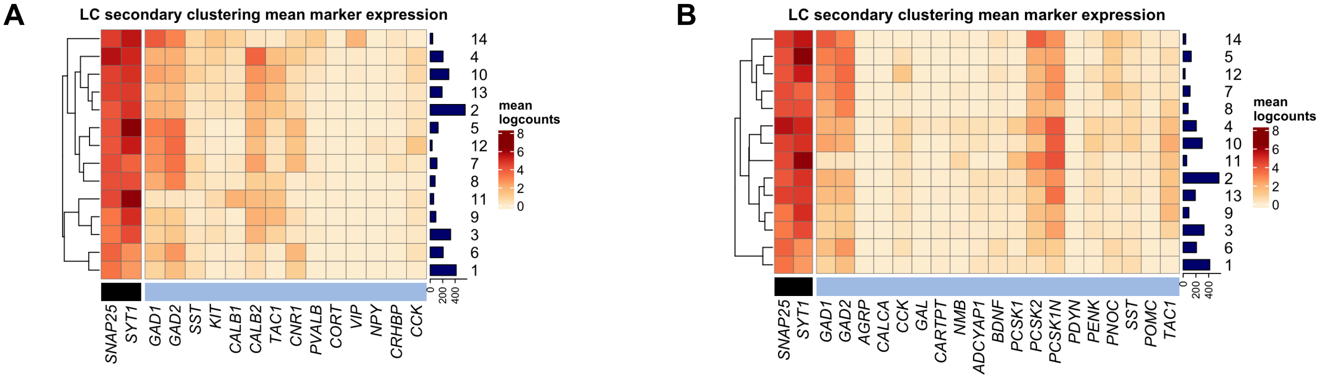


**Supplementary Figure 22: Inhibitory neuronal subpopulations identified by secondary unsupervised clustering on inhibitory neurons in snRNA-seq data.** We applied a secondary round of unsupervised clustering to the inhibitory neuron nuclei identified in the first round of clustering. This identified 14 clusters representing inhibitory neuronal subpopulations. **(A)** Expression of neuron marker genes (black) and inhibitory neuron marker genes (light blue) (columns) in the 14 clusters (rows). **(B)** Expression of neuron marker genes (black) and additional set of GABAergic neuron marker genes from a recent publication using snRNA-seq in mice [32] (light blue) (columns) in the 14 clusters (rows). Cluster IDs are shown in labels on the right, and numbers of nuclei per cluster are shown in horizontal bars on the right. Heatmap values represent mean log-transformed normalized counts (logcounts) per cluster.

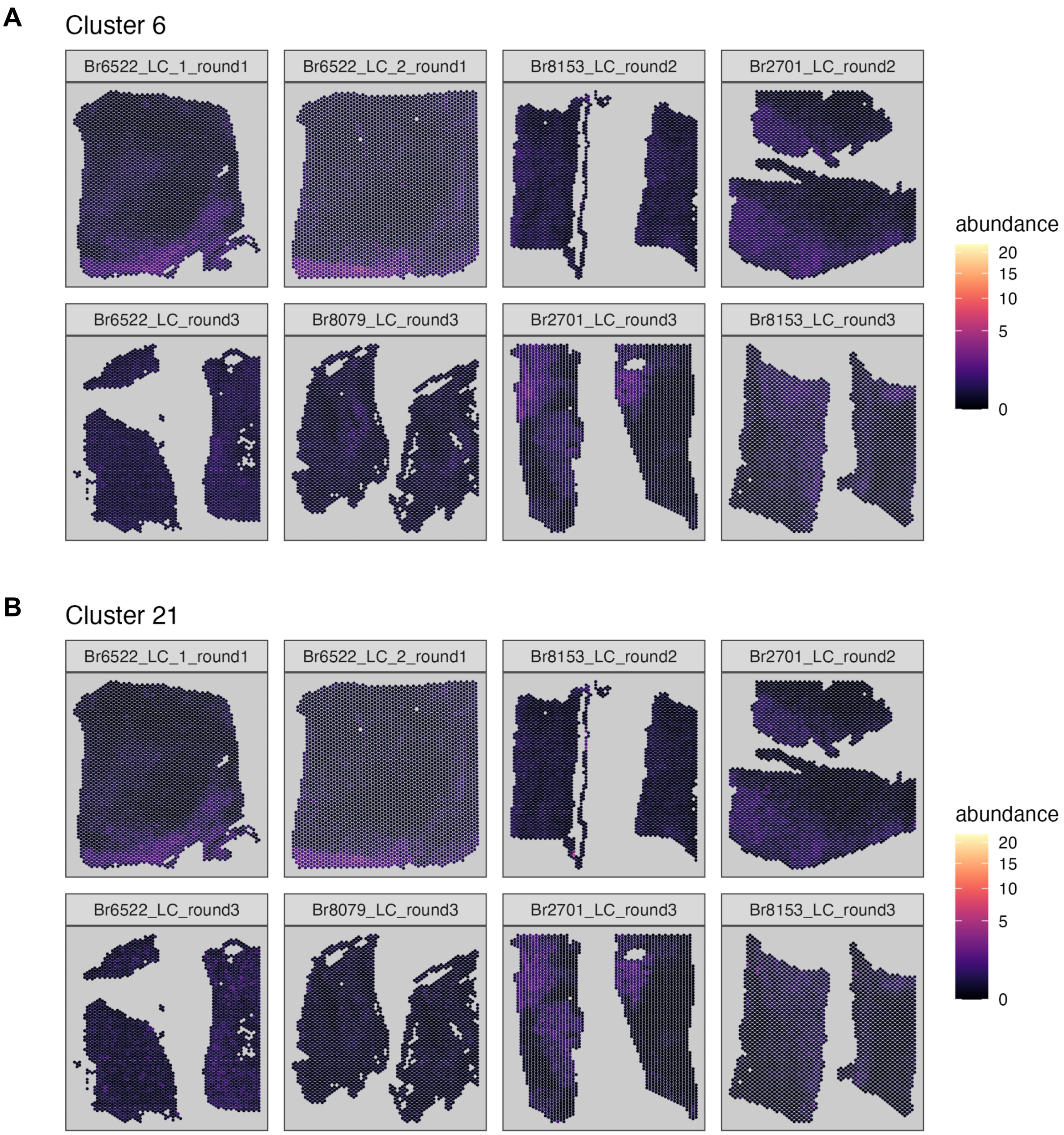


**Supplementary Figure 23: Spot-level deconvolution to map the spatial coordinates of snRNA-seq populations within the Visium SRT samples.** We applied a spot-level deconvolution algorithm (cell2location [46]) to integrate the snRNA-seq and SRT data by estimating the cell abundance of the snRNA-seq populations, which are used as reference populations, at each spatial location (spot) in the Visium SRT samples. While this approach mapped **(A)** NE neurons (cluster 6) and **(B)** 5-HT neurons (cluster 21) to the spatial regions where these populations were previously identified based on expression of marker genes (**Supplementary Figures 3** and **18**), the overall mapping performance was relatively poor. We note that these are relatively rare populations, with relatively subtle expression differences compared to other neuronal populations, and NE neurons are characterized by large size and high transcriptional activity, which may have affected performance of the algorithm. The annotated LC regions are shown in **Supplementary Figure 1**.

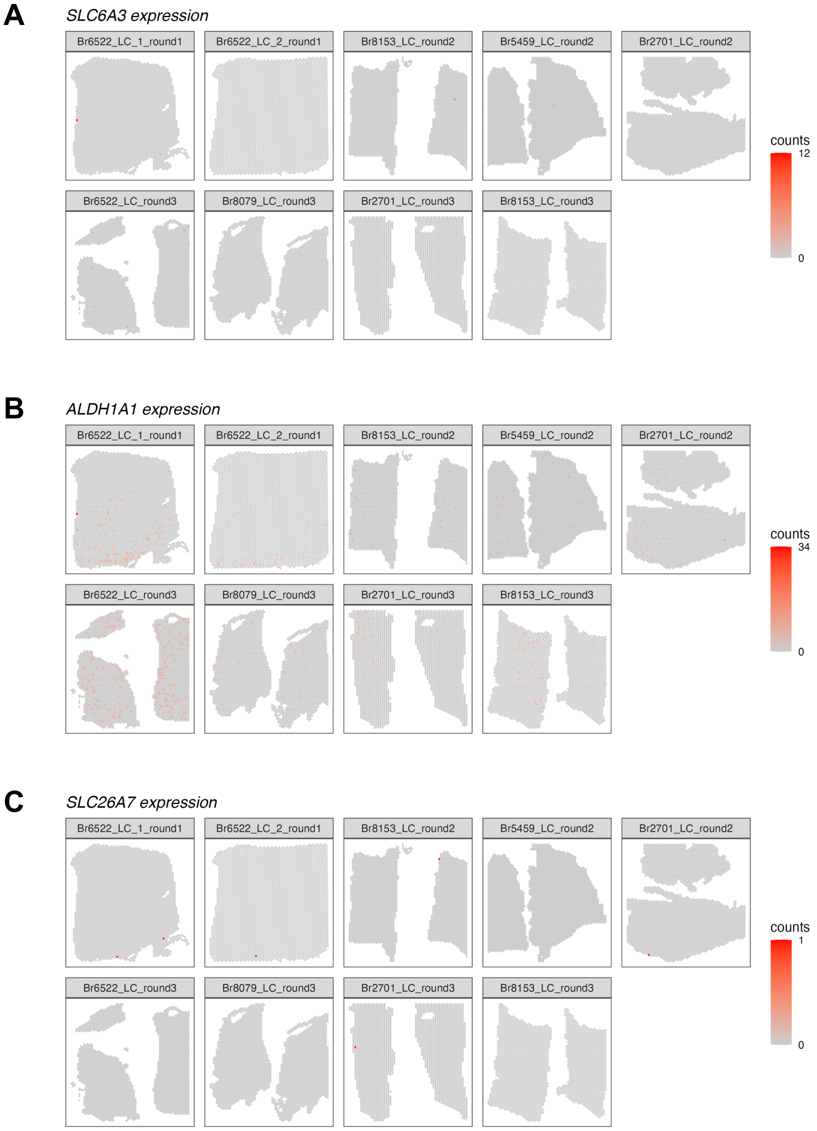


**Supplementary Figure 24: Spatial expression of dopamine (DA) neuron marker genes in Visium SRT samples.** We visualized the spatial expression of DA neuron marker genes **(A)** *SLC6A3* (encoding the dopamine transporter), **(B)** *ALDH1A1*, and **(C)** *SLC26A7* in the *N*=9 initial Visium SRT samples, which showed that these genes were not strongly expressed within the annotated LC regions. Color scale shows UMI counts per spot. The annotated LC regions are shown in **Supplementary Figure 1**.

A

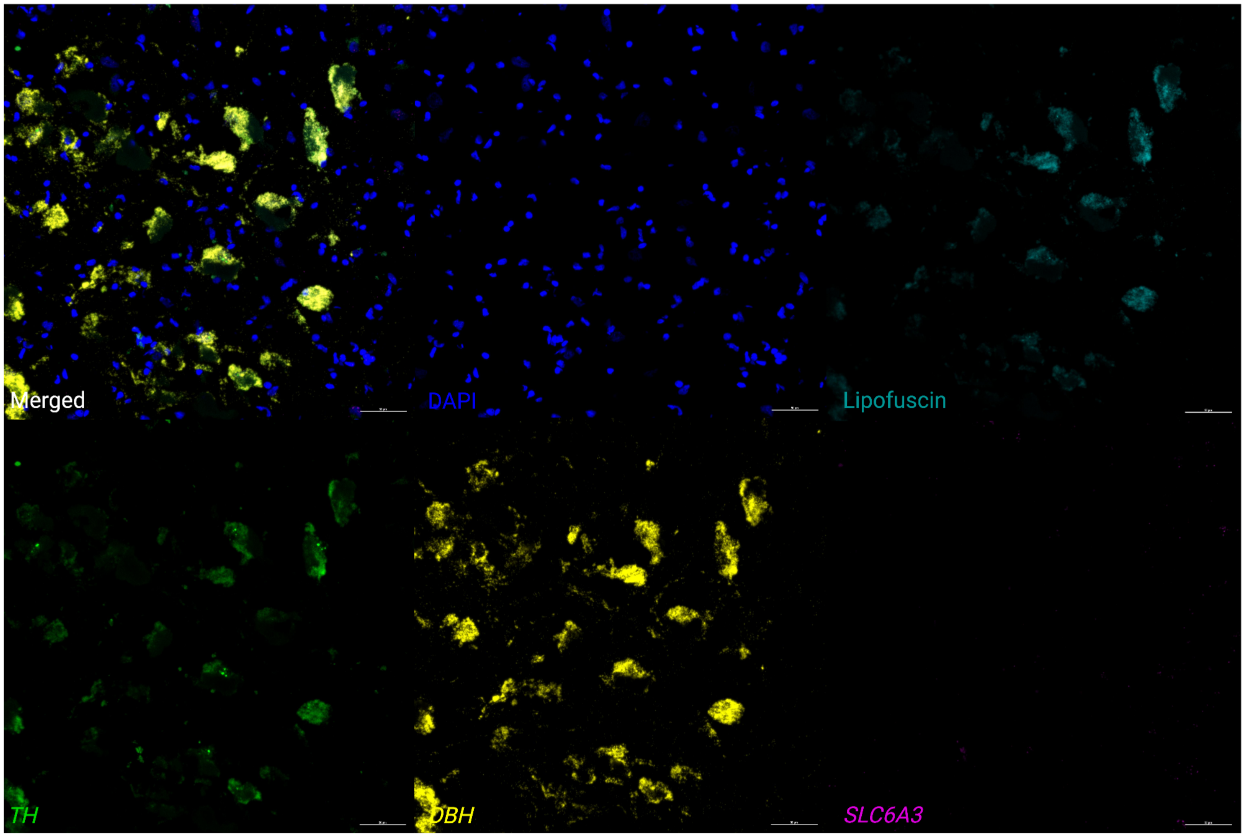
Merged
DAPI
Lipofuscin
TH
DBH
SLC6A3

B

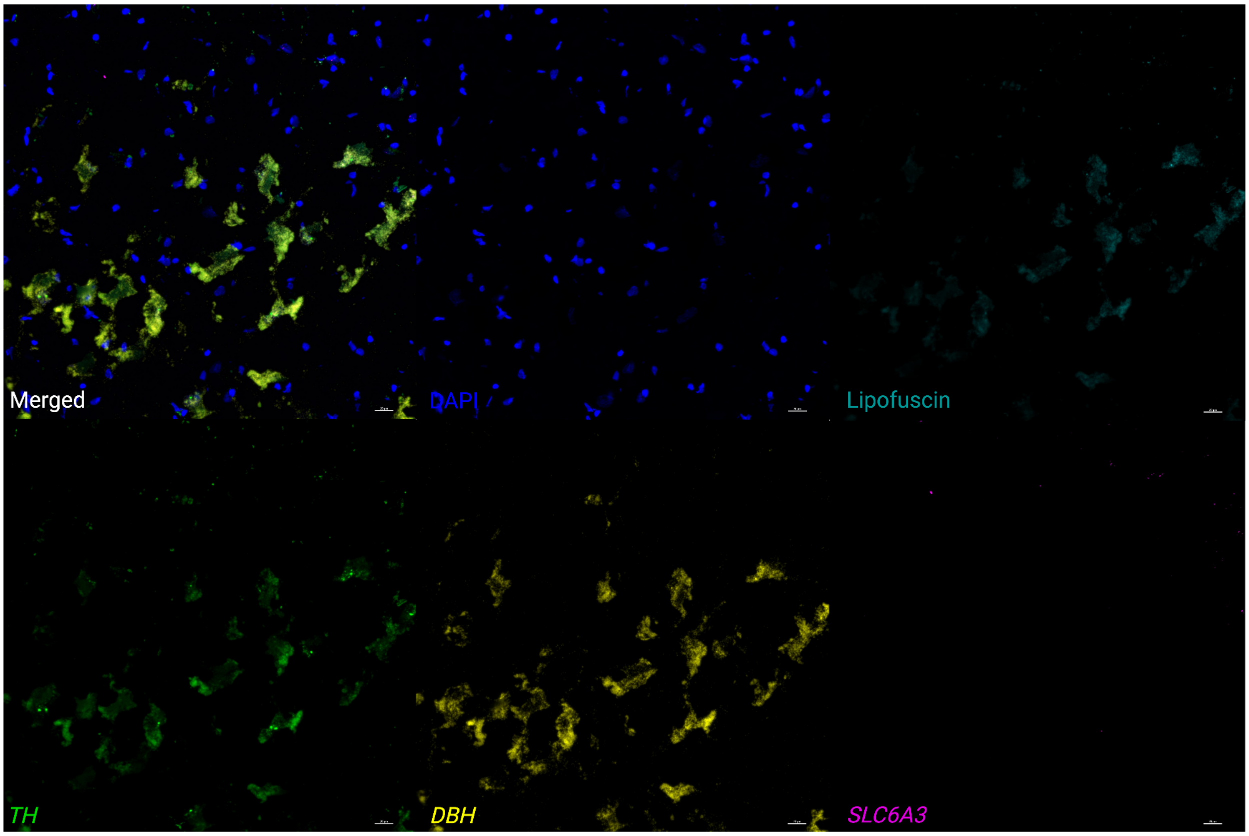
Merged
DAPI
Lipofuscin
TH
DBH
SLC6A3

**C**

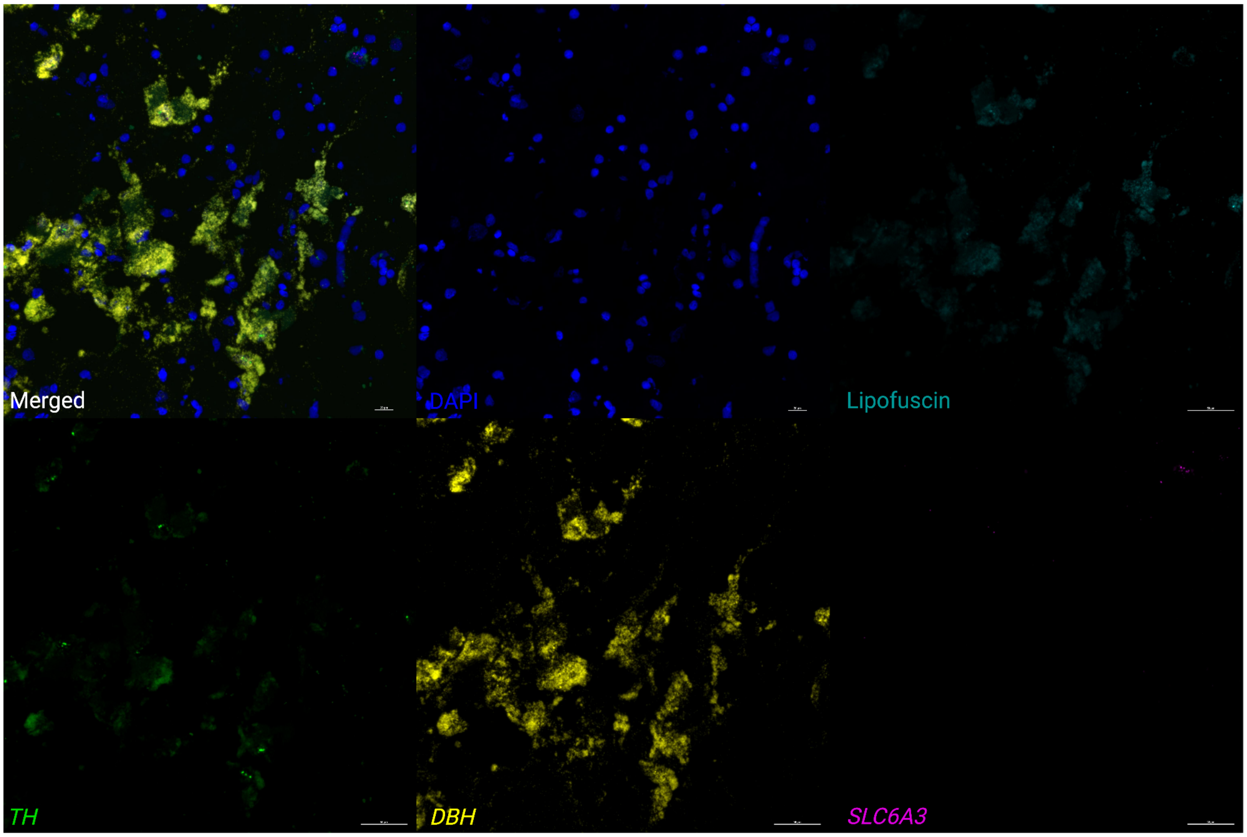


**Supplementary Figure 25: Expression of NE neuron marker genes, 5-HT neuron marker gene, and DA neuron marker gene in individual cells using RNAscope and high-magnification confocal imaging.** We applied RNAscope [37] and high-magnification confocal imaging (at 40x magnification) to samples from 3 independent donors, **(A)** Br8689, **(B)** Br5529, and **(C)** Br5426, to visualize expression of two NE neuron marker genes (*DBH* and *TH*), one 5-HT neuron marker gene (*TPH2*), and one DA neuron marker gene (*SLC6A3*) within individual cells within the LC region in each sample. We do not observe expression of the DA neuron marker gene (*SLC6A3*, encoding the dopamine transporter) within individual NE neurons (identified by co-expression of *DBH* and *TH*). Scale bar: 40 μm.

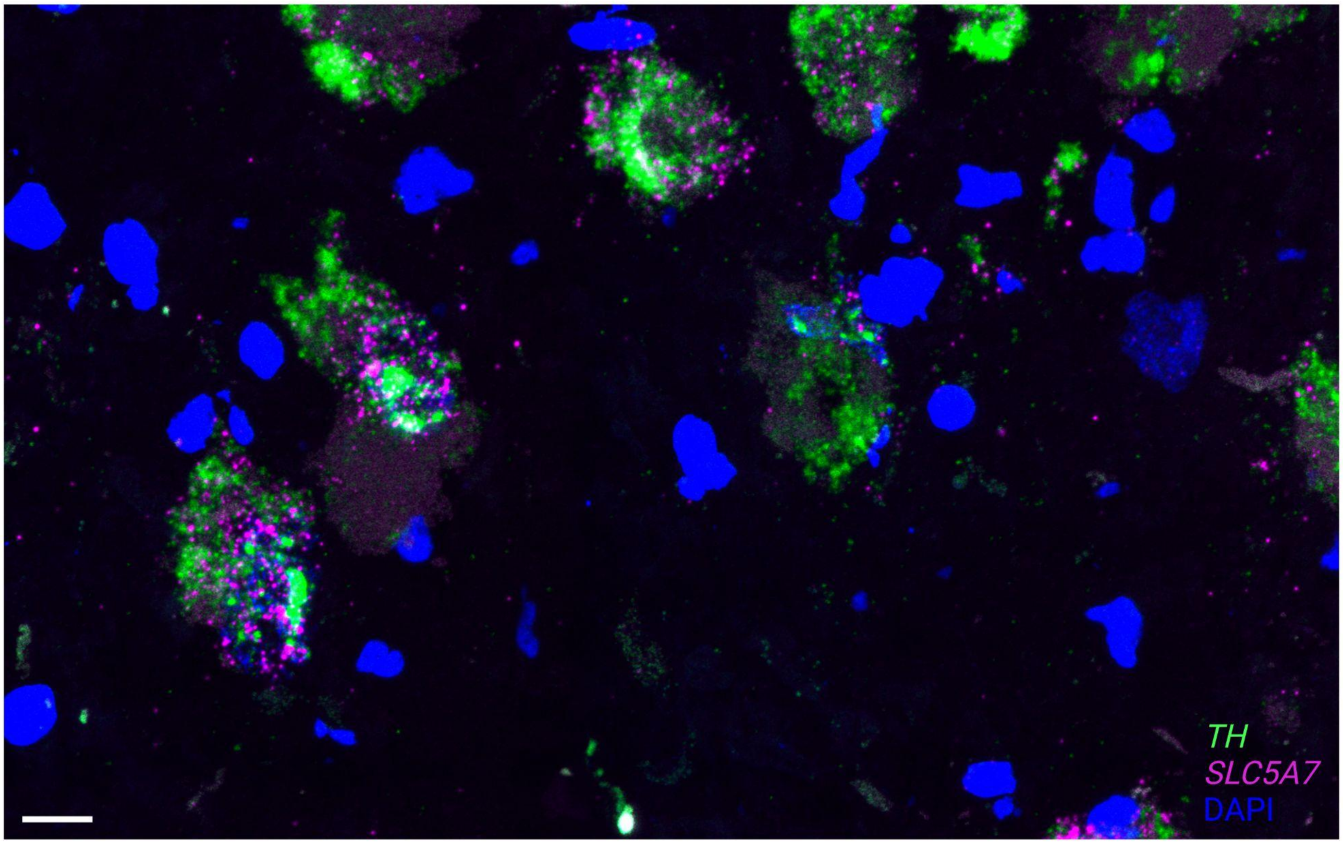


**Supplementary Figure 26: High-resolution images demonstrating co-expression of cholinergic marker gene within NE neurons.** We applied RNAscope [37] and high-resolution imaging at 63x magnification to visualize expression of *SLC5A7* (cholinergic marker gene encoding the high affinity choline transporter, shown in pink) and *TH* (NE neuron marker gene encoding tyrosine hydroxylase, shown in green), and DAPI stain for nuclei (blue), within the LC region in a tissue section from donor Br8079. This confirmed co-expression of *SLC5A7* and *TH* within individual cells. Scale bar: 25 μm.

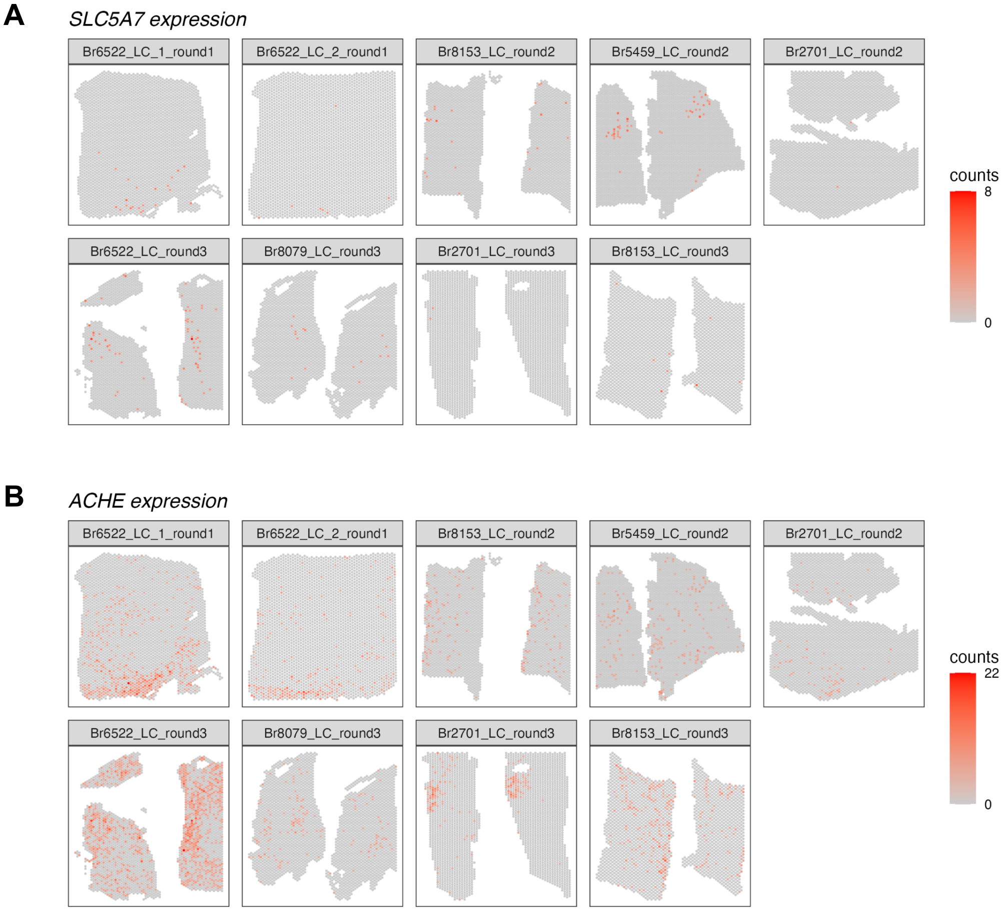


**Supplementary Figure 27: Spatial expression of cholinergic marker genes in Visium SRT samples.** We visualized the spatial expression of cholinergic marker genes **(A)** *SLC5A7* and **(B)** *ACHE* in the *N*=9 initial Visium SRT samples, which showed that these genes were expressed both within and outside the annotated LC regions. Color scale shows UMI counts per spot. The annotated LC regions are shown in **Supplementary Figure 1**.

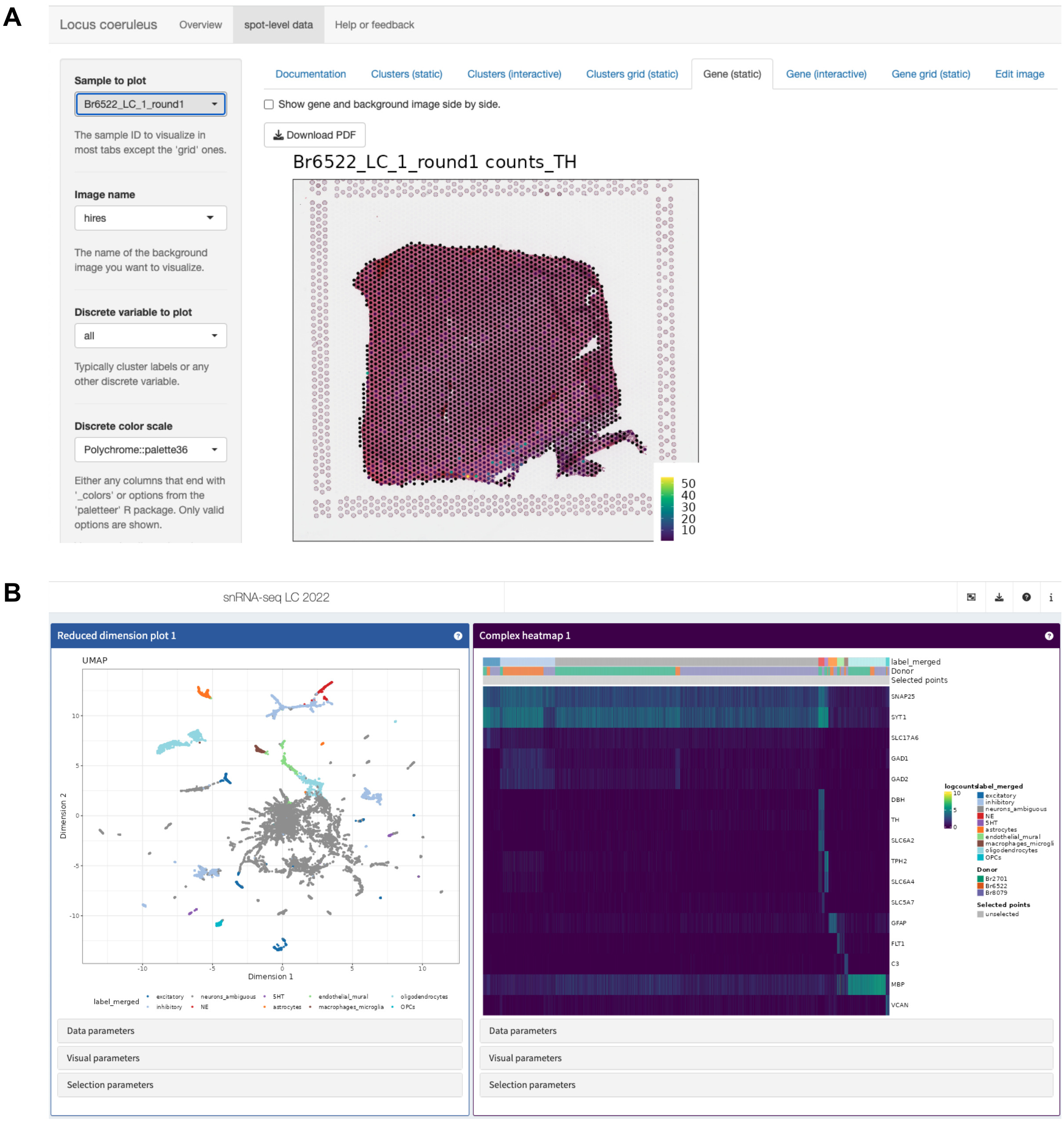


**Supplementary Figure 28: Overview of interactive web-accessible data resources.** All datasets described in this manuscript are freely accessible via interactive web apps and downloadable R/Bioconductor objects (see **Table 1** for details). **(A)** Screenshot of Shiny [50] web app providing interactive access to Visium SRT data. **(B)** Screenshot of iSEE [51] web app providing interactive access to snRNA-seq data. For instructions on how to use the web apps to search for and display individual genes, see **Supplementary Figures 29-30**.

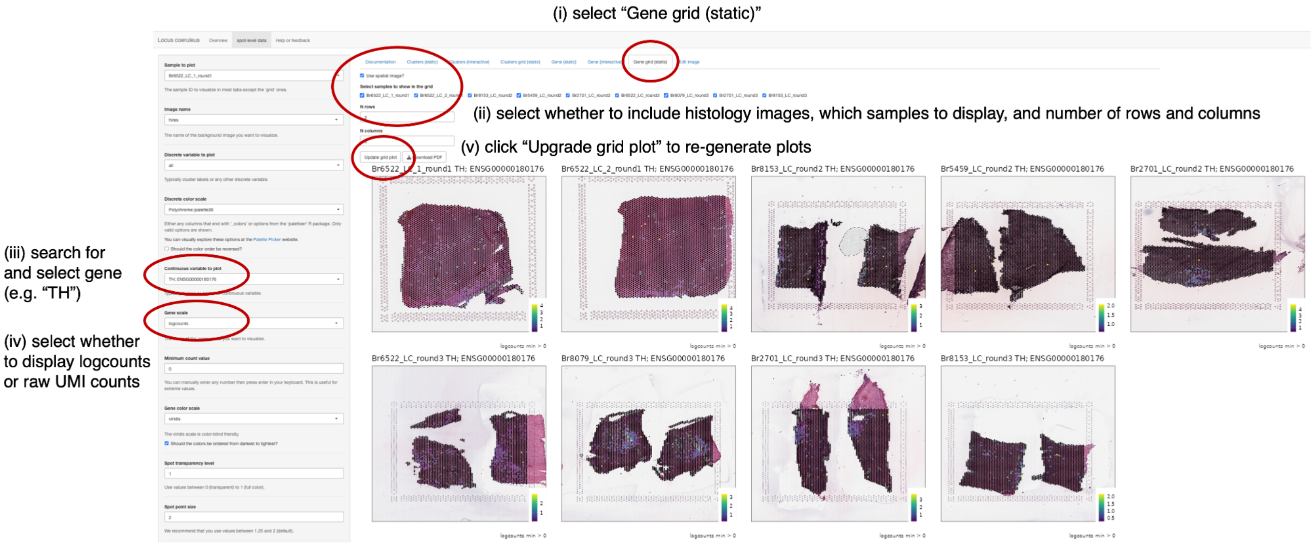


**Supplementary Figure 29: Instructions to display individual genes in Visium SRT data app.** Example displaying screenshots and sequential instructions for how to search for individual genes and show spatial gene expression in Shiny [50] web app providing interactive access to Visium SRT data.

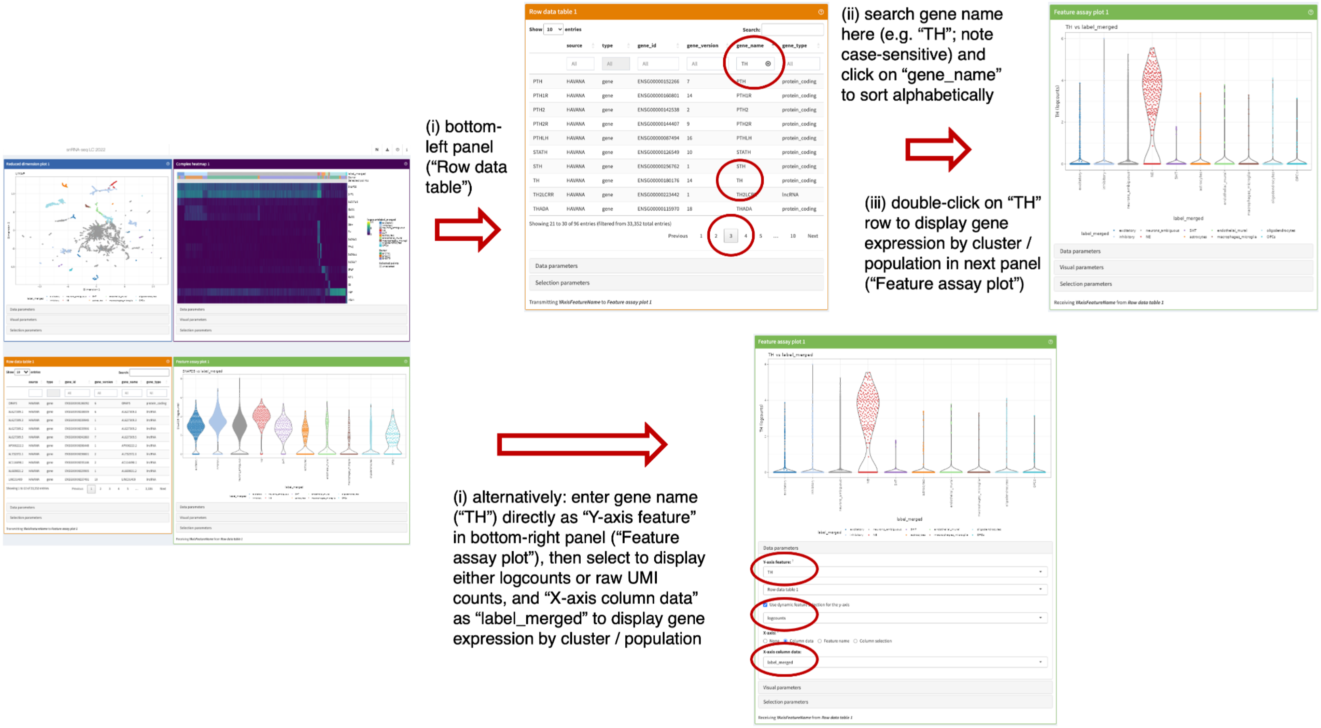


**Supplementary Figure 30: Instructions to display individual genes in snRNA-seq data app.** Example displaying screenshots and sequential instructions for how to search for individual genes and show gene expression by cluster or cell population in iSEE [51] web app providing interactive access to snRNA-seq data.
